# CopyMix: Mixture Model Based Single-Cell Clustering and Copy Number Profiling using Variational Inference

**DOI:** 10.1101/2020.01.29.926022

**Authors:** Negar Safinianaini, Camila P. E. de Souza, Andrew Roth, Hazal Koptagel, Hosein Toosi, Jens Lagergren

**Affiliations:** School of Electrical Engineering and Computer Science, KTH Royal Institute of Technology, Stockholm, Sweden; Department of Statistical and Actuarial Sciences, University of Western Ontario, London, ON, Canada; Science for Life Laboratory, Solna, Sweden; Department of Computer Science, University of British Columbia, Vancouver, BC, Canada; British Columbia Cancer Agency, Vancouver, BC, Canada

**Keywords:** Mixture Models, Variational Inference, Copy number profiling, Single-cell

## Abstract

Investigating tumor heterogeneity using single-cell sequencing technologies is imperative to understand how tumors evolve since each cell subpopulation harbors a unique set of genomic features that yields a unique phenotype, which is bound to have clinical relevance. Clustering of cells based on copy number data obtained from single-cell DNA sequencing provides an opportunity to identify different tumor cell subpopulations. Accordingly, computational methods have emerged for single-cell copy number profiling and clustering; however, these two tasks have been handled sequentially by applying various ad-hoc pre- and post-processing steps; hence, a procedure vulnerable to introducing clustering artifacts. Moreover, clonal copy number profiling has been missing except for one method, CONET, which unfortunately computes it by a post-processing tool. Finally, a common copy number profiling tool, HMMcopy, requires parameter tuning. We avoid the clustering artifact issues and provide clonal copy number profiles without the labor of parameter tuning in our method, CopyMix, a Variational Inference for a novel mixture model, by jointly inferring cell clusters and their underlying copy number profile. We evaluate CopyMix using simulated data and published biological data from ovarian cancer. The results reveal high clustering performance and low errors in copy number profiling. These favorable results indicate a considerable potential to obtain clinical impact by using CopyMix in studies of cancer tumor heterogeneity.

## 1 Introduction

A tumor typically consists of a collection of heterogeneous cell populations, each having distinct genetic and phenotypic properties, in particular, concerning the capacity to promote cancer progression, metastasis, and therapy resistance Eirew et al. (2015); Nowell (1976). Single-cell sequencing technologies Gawad et al. (2016); Navin et al. (2011); Shapiro et al. (2013); Zahn et al. (2017) provide an opportunity to investigate the genomic profile of individual cells regarding both single nucleotide variation (SNV) and copy number variation (CNV). CNVs and SNVs are essential contributors to phenotypic variation relating to health, and disease Baslan et al. (2012); Lawson et al. (2018). Although single-cell SNV profiling is hampered by experimental imperfections such as drop-outs, copy number profiling, i.e., detecting single-cell CNVs, is feasible, at least at coarser resolutions. Clustering cells based on their copy number profiles improves understanding of tumor subpopulations and tumor heterogeneity, issues bound to have clinical relevance.

Current single-cell datasets pose a wealth of computational challenges. As answers to some of those, methods have emerged for single-cell copy number profiling and clustering; some methods infer clustering after copy number profiling, e.g., Garvin et al. (2015); Zahn et al. (2017); Leung et al. (2017); Vitak et al. (2017); Zaccaria and Raphael (2021), while a recent method, CONET Markowska et al. (2022), derives copy number profiles by post-processing in addition to a user-defined sequence of breakpoints per cell. Unfortunately, these methods perform single-cell copy number profiling and clustering sequentially with various ad-hoc processes. The sequential approach is vulnerable to artifacts since preprocessing decisions, typically irreversible, constrain all later analyses. Even if each task performs optimally, the final result may still fall short of the best possible performance Blocker and Meng (2013). Another problem is when using HMMcopy Shah et al. (2006); Vitak et al. (2017); Zahn et al. (2017) for copy number profiling— copy number profiles can naturally be modeled by a sequence of latent variables forming Hidden Markov Models (HMMs)—that has limitations such as requiring manual calibration of more than ten parameters Mallory et al. (2020b). Moreover, as addressed recently by CONET Markowska et al. (2022), one should account for the fact that copy number profiles are generated by a clonal process; however, mixtures of Hidden Markov Models (MHMMs) Smyth (1997), a joint inference alternative for performing the two biological tasks, consider a different copy number profile per cell, thus lacking the clonal copy number profiling.

By a joint inference solution, also motivated by future directions of computational modeling in single-cell cancer genomics Zhang and Campbell (2020), we propose a novel framework referred to as CopyMix to alleviate three problems: (1) the sequential treatment of copy number profiling and clustering; (2) the labor of HMMcopy parameter tuning; (3) lack of clonal copy number profiling by MHMMs. CopyMix performs the joint inference of copy number profiling and clustering, and as to the best of our knowledge, no earlier work has simultaneously performed these two tasks. Similarly, joint inference has been considered for single-cell DNA methylation Kapourani and Sanguinetti (2019); de Souza et al. (2020), single-cell SNV data Roth et al. (2016), and bulk chIP-seq data from several replicates Zuo et al. (2016). Due to the model-based treatment, CopyMix enjoys the advantages of a fully probabilistic framework, such as transparency, uncertainty measurements, and modeling flexibility. CopyMix, a biologically meaningful tangent of MHMMs, uses a novel mixture model with components (expressing the clonal process as opposed to cell-specific MHMMs) corresponding to clusters, each having a specific copy number profile, revealing the copy number variation pattern behind each cluster. We deploy a Bayesian treatment and infer all quantities of interest using Variational Inference (VI) Jordan et al. (1999), which typically yields faster inference methods than methodologies like Markov chain Monte Carlo sampling Blei et al. (2017). Compared to Expectation-Maximization (EM), VI has multiple advantages; e.g., it estimates the posterior distributions rather than point estimates, protects against over-fitting and allows for principled model selection, i.e., identifying the optimal number of mixture components Bishop (2006). Finally, our novel VI simplifies the calculations needed for the posterior approximation of the HMM part, while earlier methods McGrory and Titterington (2009b) use complex solutions to normalize the posterior approximation.

We evaluate CopyMix by experiments on biologically inspired simulated data and two published biological data Laks and McPherson (2019); Leung et al. (2017). In Laks and McPherson (2019), HMMcopy is used as a copy number profiling tool proceeding by an ad-hoc combination of clustering methods: K-means, Hierarchical Density-Based Spatial Clustering, and UMAP. We also compare CopyMix results with Ginkgo Garvin et al. (2015)—a state-of-art framework for clustering of single-cells based on previously estimated copy number profiles obtained by circular binary segmentation method Olshen et al. (2004), outperforming HMMCopy in copy number profiling although HMMCopy provides higher quality for breakpoint detection Mallory et al. (2020a). Lastly, we extend CopyMix to a so-called SNV-CopyMix for exploiting the SNV signals by incorporating SNVs in the proposed model.

A sequential version of CopyMix, resembling Shah et al. (2006); Vitak et al. (2017) but excluding their ad-hoc processes, is compared to CopyMix, confirming the hypothesis that a joint inference framework is superior over a sequential version of the framework.

## 2 CopyMix

CopyMix is a probabilistic clustering method based on a mixture of Markov chains. CopyMic is a mixture model with components corresponding to clusters, each having a specific copy number profile modeled by a sequence of latent variables. Similarly, as in Shah et al. (2006); Vitak et al. (2017); Zahn et al. (2017), we assume that each latent copy number sequence is governed by a Markov chain (describing a sequence of possible copy numbers in which the probability of each copy number value, defined as a state, depends only on the previous state in the sequence). We provide two observational models for CopyMix. The first one is based on Poisson emission, and the second one is based on Gaussian emission. In the following sections, we describe these two choices of emission modeling.

After presenting our method, CopyMix, we extend CopyMix to incorporate single nucleotide data; we refer to this method as SNV-CopyMix. The purpose of designing SNV-CopyMix is to exploit CNV-independent signals, i.e., SNVs, when performing single-cell clustering.

### 2.1 Poisson emission

We assume that the data (read counts) follow a Poisson distribution as in Witten (2011); we consider read counts per fixed equal size genomic bin as in Leung et al. (2017). Moreover, we assume the read counts are emitted from a latent sequence of copy number states. Fig. 1 illustrates an overview of CopyMix; it analyzes the input cell sequences, containing read counts per predefined genomic bin, and produces clusters of cells with their corresponding copy number sequences (profiles) explaining the clusters. Fig. 2 illustrates CopyMix graphical model where *Y* denotes the observable variables, read counts per predefined bins, *C* the latent copy number states forming a Markov chain, and *Z* the latent cell-specific cluster assignment variables. We notice in Fig. 2 that each *Y* has two levels of dependencies, which are reflections of the assumption that the Poisson distribution over the read counts depends on a latent cell-specific cluster assignment, *Z*, and a corresponding latent copy number state, *C*. Intuitively, a higher copy number should correspond to a higher read count. We incorporate this assumption in our model by defining the rate of Poisson distribution as the product of copy number state and cell-specific rate, *θ*_*n*_, see Eq. (1); this also implies that *θ*_*n*_ corresponds to the average sequencing coverage for a haploid genome. Due to this multiplicative structure for the Poisson rate, a copy number state of zero results in the lowest rate implying copy number deletion events; however, since the rate of Poisson cannot be zero, our implementation handles this by assigning a tiny number instead of zero. Our model is described in more detail in what follows, using conjugacy regarding the priors.

**Figure 1.**
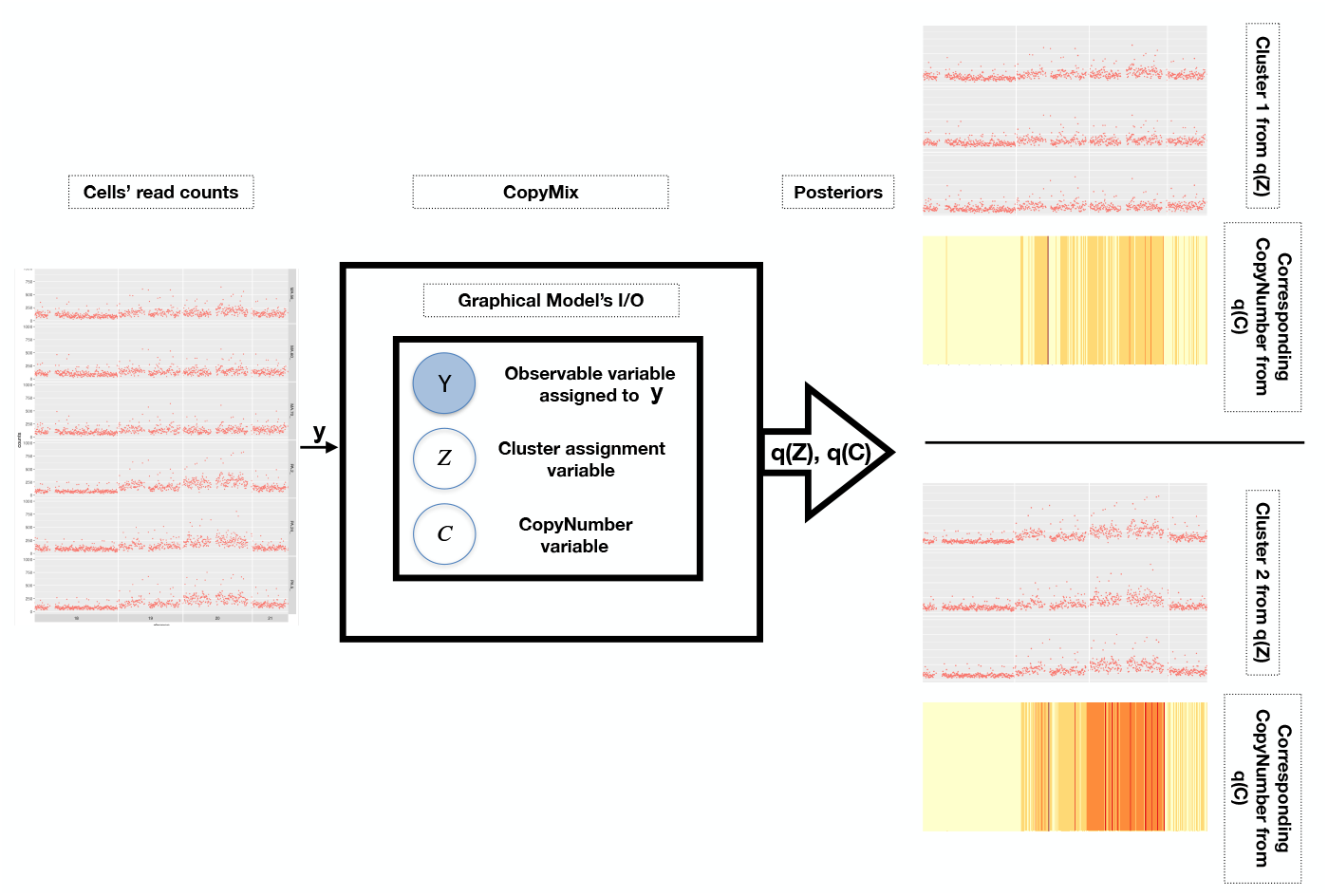
The overview of CopyMix with: input *y*, cells’ reads; outputs *q*(*Z*) and *q*(*C*), posterior distributions resulting in clusters within *y* with their corresponding copy number profiles illustrated by heatmaps (this is an example of binary clustering).

**Figure 2.**
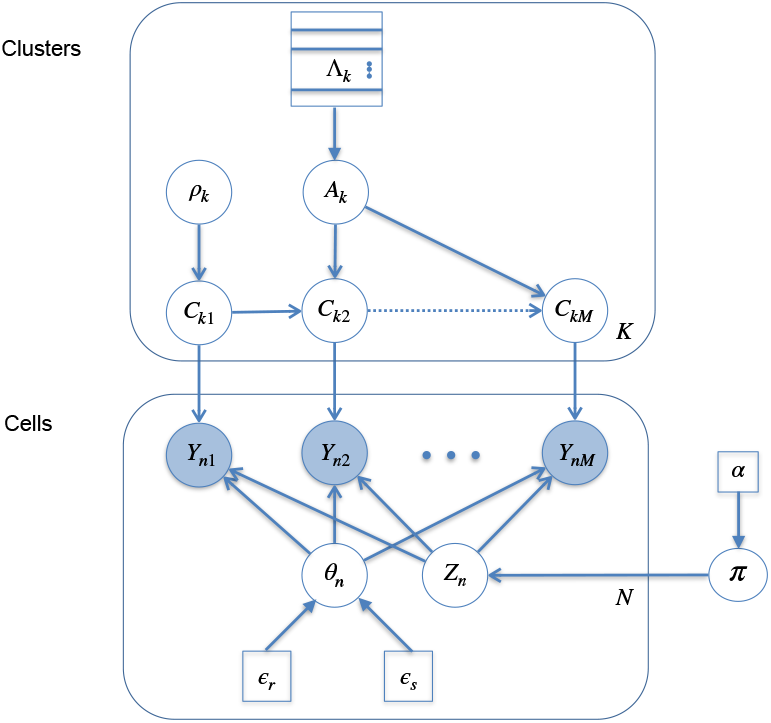
Probabilistic graphical model representing our proposed model. Shaded nodes represent observed values, the unshaded ones are the latent variables and the squares are the hyperparameters of the model; a posterior distribution over the values of the unshaded nodes is approximated using Variational Inference. *Y*_*nm*_, observed read counts from cell n and bin m; *C*_*km*_, corresponding latent copy number state forming a Markov chain; *θ*_*n*_, cell-specific rate; *Z*_*n*_, latent cell-specific cluster assignment variable. *π* and *A*_*k*_, the priors over *Z*_*n*_ and *C*_*km*_ respectively; *ρ*_*k*_, prior over the starting copy number state.

The genome considered is partitioned into *M* equally sized segments of consecutive positions called bins. Let **Y**_*n*_ = (*Y*_*n*1_, …, *Y*_*nM*_), where *Y*_*nm*_ is the random but observable number of reads aligned to bin *m* for cell *n* taking values in {0, 1, 2, …} for *m* ∈ [*M*] = {1, …, *M* } and *n* ∈ [*N*] = {1, …, *N* }. We assume that **Y**_1_,…, **Y**_*N*_ are independent and given a vector of true copy number states, *Y*_*n*1_, …, *Y*_*nM*_ are also independent with the distribution of *Y*_*nm*_ depending on the true copy number state at bin *m*. The cluster membership of cell *n* is indicated by the hidden variable *Z*_*n*_ that takes values in [*K*] = {1, …, *K*}. We assume there are *K << N* vectors of true hidden copy number states, i.e., one for each cluster. The variables *Z*_1_, …, *Z*_*N*_ are independent following a categorical distribution with *P*(*Z*_*n*_ = *k*) = *π*_*k*_ and 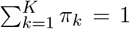. If *Z*_*n*_ = *k* then the distribution of **Y**_*n*_ depends on the *k*-th vector of true hidden copy number states, defined as **C**_*k*_ = (*C*_*k*1_, …, *C*_*kM*_), with each *C*_*km*_ taking values in [*J*] = {1, …, *J*}. We assume that *C*_*k*1_, …, *C*_*kM*_ follow a discrete-time homogeneous Markov chain with initial probabilities *ρ*_*kj*_ = *P*(*C*_*k*1_ = *j*), *j* ∈ [*J*] and transition probabilities 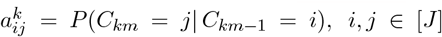. Consequently, given the cluster assignment and the corresponding true hidden vector of copy number states, *Y*_*n*1_, …, *Y*_*nM*_ are independent with *Y*_*nm*_ following a distribution with parameters depending on the hidden true state at bin *m* for cluster *k*, that is, 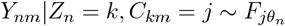. As mentioned earlier, we assume 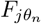 to be a Poisson distribution with rate *j × θ*_*n*_, that is,

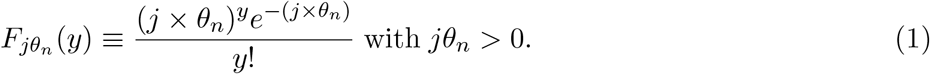

We, w.l.o.g., consider the initial probabilities, *ρ*_*kj*_’s, to be fixed and known. We let **Ψ** be the set containing all the unknown model parameters, i.e., **Ψ** = {**A, *θ, π***}, where **A** = {**A**_*k*_ : *k* ∈ [*K*]} with 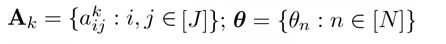; and ***π*** = (*π*_1_, …, *π*_*K*_). In order to infer **Ψ**, the hidden states **C** = {**C**_1_, …, **C**_*K*_}, and **Z** = (*Z*_1_, …, *Z*_*n*_) we apply VI; that is, we derive an algorithm that, for given data, approximates the posterior distribution of the parameters by finding the Variational Distribution (VD), *q*(**Z, C, Ψ**), with smallest Kullback-Leibler divergence to the posterior distribution *P*(**Z, C, Ψ**|**Y**), which is equivalent to maximizing the evidence lower bound (ELBO) Blei et al. (2017) given by

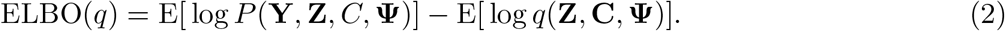

We consider the following prior distributions for the parameters in **Ψ**.

- 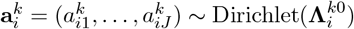.
- 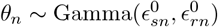, with 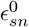, shape, and 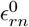, rate, hyperparameters.
- *π* ∼ Dirichlet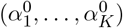

The main steps of our VI approach, for inferring **Z, C**, and **Ψ**, are described below.

#### Step 1. VD factorization

We assume the following factorization of the VD:

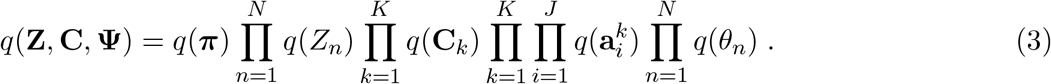

#### Step 2. Joint distribution

The logarithm of the joint distribution satisfies: log *P*(**Y, Z, C, Ψ**) = log *P*(**Y**|**Z, C, Ψ**) + log *P*(**C**|**Ψ**) + log *P*(**Z**|**Ψ**) + log *P*(**Ψ**). For details of the calculations see Appendix A.1.

#### Step 3. VD computation

We now derive a coordinate ascent algorithm for the VD. That is, we derive an update equation for each term in the factorization, Eq. (3), by calculating the expectation of log *P*(**Y, Z, C, Ψ**) over the VD of all random variables except the one currently being updated Bishop (2006). For example, we obtain the update equation for ***π***, *q*(***π***), by calculating *E*_−_***π***( log *P*(**Y, Z, C, Ψ**)), where −***π*** indicates that the expectation is taken with respect to the VD of all other random variables than ***π***, i.e., **Z, C** and {**A, *θ***} except ***π***. See A.2 and A.3 for the update equation of each term in Eq. (3).

#### Step 4. Summary of updates

We update the parameters of each posterior distribution presented in Step 3; for the summary of the update equations, see Alg. 2, in Appendix A.2. A shorter version of Alg. 2, providing an overview of CopyMix, is shown in Alg. 1. As shown in lines 8 and 9 of Alg. 1, we aim to find the posterior distributions of cluster assignments and copy number states. Note that the update equations in line 5 of Alg. 1 can be run in a parallel fashion, keeping the algorithm’s computational complexity low. The implementation of CopyMix is available at **CopyMix**.

##### Algorithm 1 CopyMix: a Variational Inference Algorithm

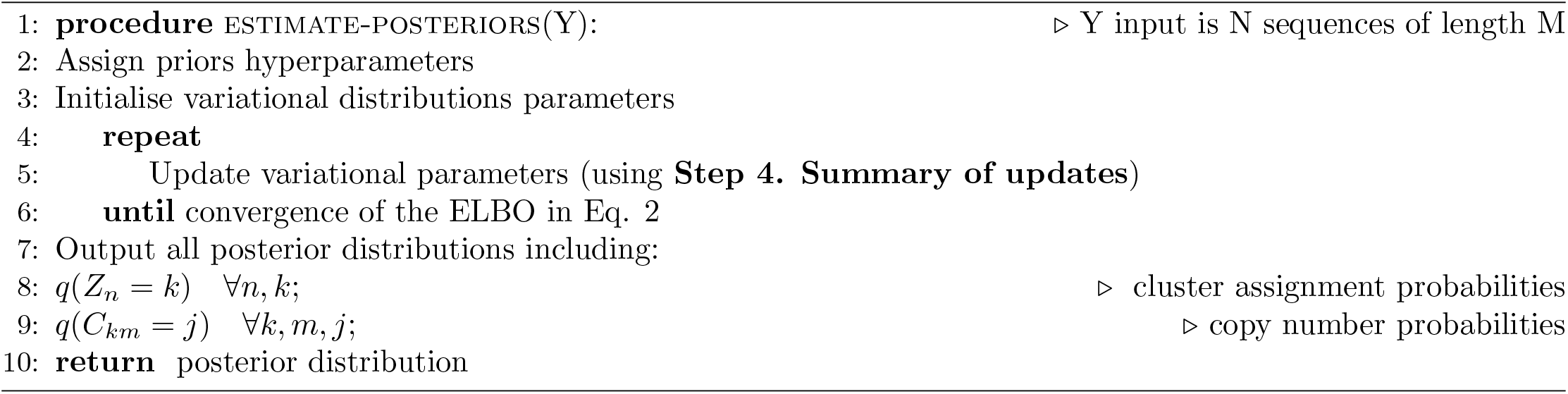

### 2.2 Gaussian emission

In this version, our input data, read count ratios (or read ratios), are assumed to be GC-corrected—a necessary bias correction due to the dropping of read coverage at the regions with extreme GC contents Yoon et al. (2009). The read counts are integers, and the GC-corrected read counts become positive real numbers (read ratios) through the GC-correction process. We consider read ratios over *M* fixed, typically equal-sized, genomic bins, as in Leung et al. (2017); Laks and McPherson (2019). We assume that the read ratios follow a Gaussian distribution; for the details supporting this choice, see Appendix A.10. Moreover, we presume that the read ratios are emitted from a latent sequence of copy number states. The probabilistic graphical model is illustrated in Fig. 3. The read ratios are modeled as a Gaussian distribution with independent conjugate priors; that is, the mean of the Gaussian follows a Gaussian distribution and the precision of the Gaussian follows a Gamma distribution. *Y* denotes the observable variables, read ratios per predefined bins, *C* the latent copy number states forming a Markov chain, and *Z* the latent cell-specific cluster assignment variables. Note that in the Results section, we account for chromosomal copy numbers by assigning initial probabilities to the beginning of each chromosome.

**Figure 3.**
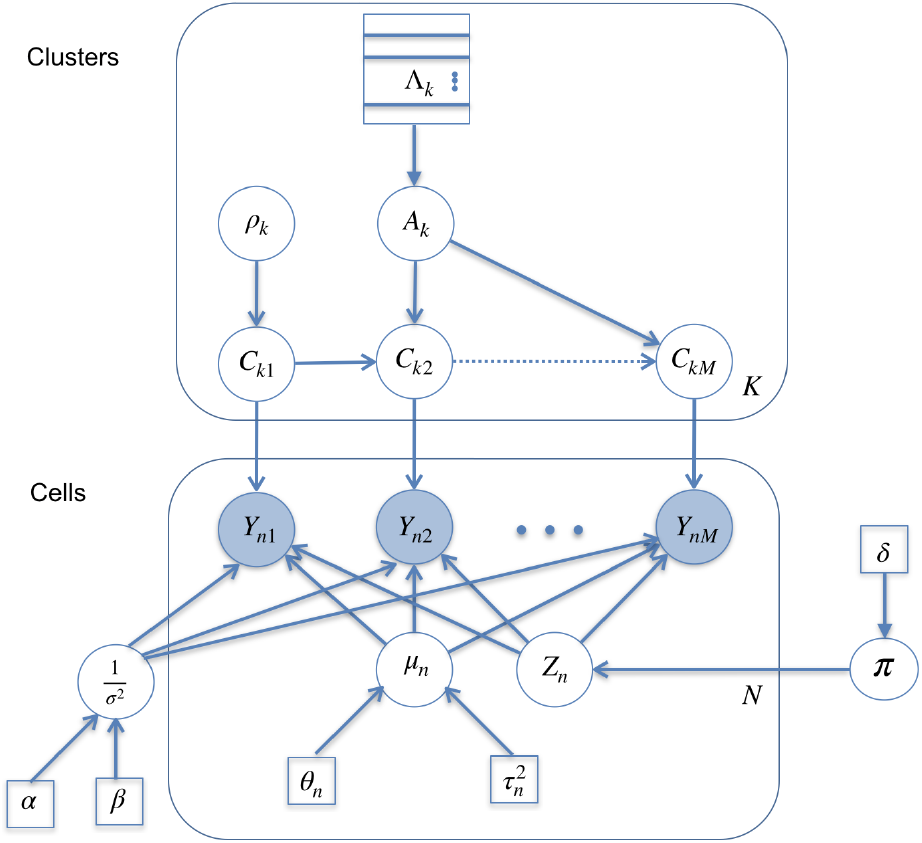
In this version *Y*_*nm*_ are observed corrected read ratios from cell n and bin m; C_*km*_, corresponding latent copy number state forming a Markov chain; *μ*_*n*_, cell-specific rate; *Z*_*n*_, latent cell-specific cluster assignment variable. *π* and *A*_*k*_, the priors over *Z*_*n*_ and *C*_*km*_ respectively; *ρ*_*k*_, prior over the starting copy number state.

Let *Y*_*nm*_ be the observed gc-corrected reads ratio to bin *m* for cell *n* for *n* = 1, …, *N* and *m* = 1, …, *M*. Let **Y**_*n*_ = (*Y*_*n*1_, …, *Y*_*nM*_) be vector with all observed data for cell *n*. Given the cluster assignment and the corresponding true hidden vector of copy number states, *Y*_*n*1_, …, *Y*_*nM*_ are independent with *Y*_*nm*_ following a distribution with parameters depending on the hidden true state at bin *m* for cluster population *k*, that is,

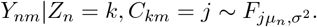

We assume 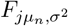 to be a Gaussian distribution with *jμ*_*n*_ = *j × μ*_*n*_, i.e.,

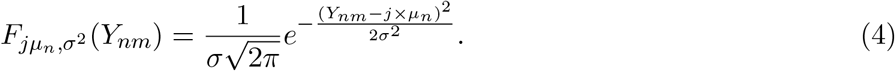

The use of the multiplicative structure in the mean of the Gaussian distribution can be found in a recent work in which a distribution mean is dependent on copy number events by a multiplication operation Malekpour et al. (2018). Note that one needs B-allele frequencies (BAF) to perform ploidy estimation, and the methods not using BAF data apply restrictive assumptions to select among many equally plausible solutions to estimated ploidy Zaccaria and Raphael (2021); this may result in selecting copy numbers that contradict the underlying allelic balance/imbalance. In the Concluding remarks, we refer to allele-based CopyMix as a future work.

Let **Ψ** be the set containing all the unknown model parameters, i.e., **Ψ** = {**A, *μ***, *σ*^2^, ***π***}, where

- **A** = {**A**_*k*_ for *k* = 1, …, *K*} with 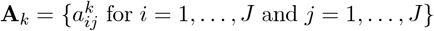;
- ***μ*** = {*μ*_*n*_ for *n* = 1, …, *N* };
- *σ*^2^;
- ***π*** = (*π*_1_, …, *π*_*K*_).

We consider the following prior distributions for the parameters in **Ψ**.

- 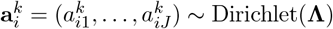
- 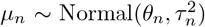. The conjugate prior concerning the mean of Normal distribution is Normal distribution.
- 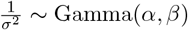. The conjugate prior concerning the precision of Normal distribution is Gamma distribution.
- ***π*** ∼ Dirichlet(***δ***)

The main steps of our VI approach, for inferring **Z, C**, and **Ψ**, are described below.

#### Step 1. VD factorization

We assume the following factorization of the VD:

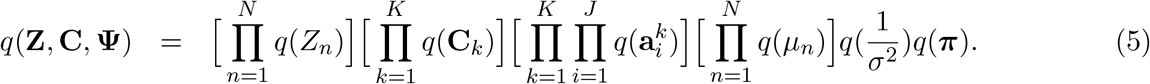

#### Step 2. Joint distribution

The logarithm of the joint distribution satisfies: log *P*(**Y, Z, C, Ψ**) = log *P*(**Y**|**Z, C, Ψ**) + log *P*(**C**|**Ψ**) + log *P*(**Z**|**Ψ**) + log *P*(**Ψ**). For details of the calculations see Appendix A.4.

#### Step 3. VD computation

We now derive a coordinate ascent algorithm for the VD; as we described this for the Poisson version, we derive an update equation for each term in the factorization, Eq. (5), by calculating the expectation of log *P*(**Y, Z, C, Ψ**) over the VD of all random variables except the one currently being updated Bishop (2006). See A.5 and A.6 for the update equation of each term in Eq. (5).

#### Step 4. Summary of updates

We update the parameters of each posterior distribution presented in Step 3; for the details of the update equations, see Appendix A.5. We summarize the updates as shown in Alg. 1.

### 2.3 SNV-CopyMix

In this section, we incorporate SNV data into the model (see Fig. 4). We augment CopyMix graphical model by a copy-number-independent SNV process; we refer to this version of CopyMix as SNV-CopyMix. In Fig. 4, the new components are colored in magenta. *X*_*nl*_ denotes the observable variable, corresponding to a nucleotide in the genome, that we assume to be dependent on cluster-specific latent point mutation *S*_*kl*_. Naturally, each *S*_*kl*_ is distributed as Bernoulli, where mutation corresponds to value of zero, and non-mutation corresponds to value of one. The prior distribution of *S*_*kl*_ is *ξ*, a Beta distribution. Note that the SNV data contains *L* sites such that *L >> M*. In the following subsection, the calculation of the probability of *X*_*nl*_ is described; for more detail see the work by Koptagel et al. (2018). The VI update equations are calculated analogously to those in CopyMix; see A.7.

**Figure 4.**
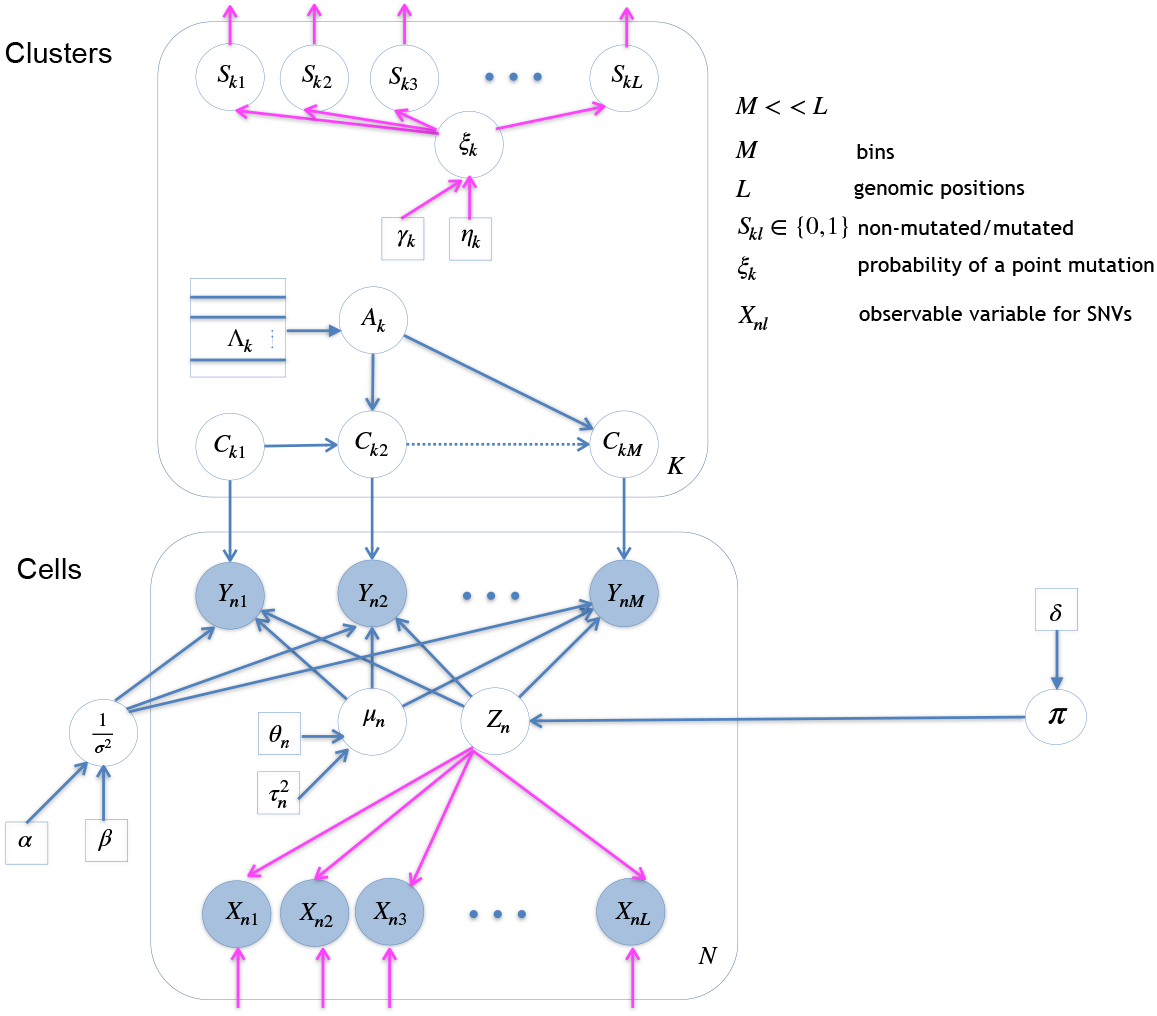
The graphical model with SNV process included.

#### 2.3.1 The likelihood model for SNVs

Here, we briefly describe the computations related to the likelihood model for the SNVs, i.e., the probability of each *X*_*nl*_ is calculated. For more details, see the earlier work Koptagel et al. (2018). The genotype of a site *l* is denoted by 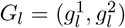 where 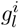 is the nucleotide of allele *i*. The site genotype consists of two copies of the reference nucleotide in the case of no mutation. It consists of one reference (denoted as *ref*) and one alternate (denoted as *alt*) nucleotide in the case of a heterozygous SNV, and two alternates in the case of homozygous SNV. Note that the sites are assumed to be diploid. Let *t* be an arbitrary nucleotide in the read. Also, assume that *r* is an enumeration of the number of reads; e.g., if a cell has five reads at a site, then *r* = 1, .., 5. Moreover, we define *ϵ* to be a the error probability. It is computed by processing the Phred scores in data—Phred score Ewing et al. (1998) is a measure of the quality of the identification of the nucleotides generated by DNA sequencing. The calculation of genotype likelihood is similar to Monovar Zafar et al. (2014), but it is a generalized version of Monovar, where we explicitly distinguish allele info for *refref* and *altalt* cases. To this end, the likelihood of a single read of cell *n* at site 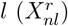 with the corresponding error probability 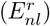 is

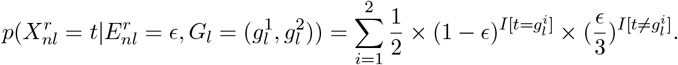

The reads are assumed to be i.i.d., so the likelihood of the reads of cell *n* at site *l* is

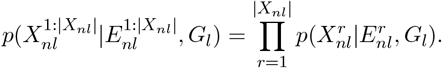

The likelihood for a non-mutation case is 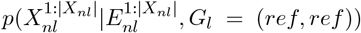. The likelihood for a heterozygous mutation case is 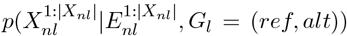, and a homozygous mutation case is 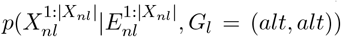. Note we allow for two versions of SNV-CopyMix, that is, (1) likelihood conditioned on non-mutation versus homozygous mutation, and (2) likelihood conditioned on non-mutation versus heterozygous mutation. In the experiment section, we test both of these versions when referring to SNV-CopyMix.

## 3. Empirical investigation

### 3.1 Experimental setup

#### 3.1.1 Dataset

To adequately evaluate CopyMix, as opposed to earlier literature lacking the evaluation against ground truth and hence subjective, we conduct experiments on simulated data, where we have access to the ground truth. We also evaluate CopyMix on two biological datasets Laks and McPherson (2019); Leung et al. (2017): (1) Read count data from Leung et al. (2017) (available from the NCBI Sequence Read Archive (SRA) under accession number SRP074289); (2) Read ratio data from Laks and McPherson (2019) accessed via DLP.

The first biological data stem from single-cell whole-genome sequencing from 18 primary colon cancer tumor cells and 18 metastatic cells from matched liver samples for one patient referred to as CRC2 in Leung et al. (2017). In this work, we mainly analyze CRC2 patient data (more complex than CRC1 according to Leung et al. (2017)) considering chromosomes 18, 19, 20, and 21 comprising a total of 904 genomic bins of an approximate size of 200Kb each. Each data point corresponds to the number of reads aligned per bin per cell after GC correction Leung et al. (2017). These counts stemming from the tumor cell’s genome reflect the hidden copy numbers—copy number integers are conventionally ranged from 0 to 6, as in Leung et al. (2017). The higher the counts are, the higher the copy numbers are expected to be. Regarding clustering in Leung et al. (2017), it is reasonable that the primary tumor cells from the colon and the metastatic cells from the liver should cluster separately, as inter-tumor copy number differences are expected to dominate over minor intra-tumor copy number variations; therefore, we use the terms primary and metastatic clusters.

The second biological data stems from single-cell whole-genome sequencing using DLP technology. The data is from three cell lines from ovarian cancer with a total number of 891 cells Laks and McPherson (2019). It is meaningful to infer that each cell line is a separate cluster; however, the goal is to find sub-clusters as well.

For the simulated data, we examine both Poisson and Gaussian models. For the Poisson version, we randomly generate 30 datasets for each test case, following our graphical model and inspired by biological datasets from Leung et al. (2017); for data simulation details, see A.8. We examine different test cases, orthogonal to each other, to show our approach’s robustness; each scenario, for both clustering and copy number state inference, is reproducible. Regarding the Gaussian model, we similarly show the robustness of CopyMix by varying the number of clusters and clustering scenarios. We simulate data under various clustering scenarios, varying the number of clusters and copy number profiles. We assume a diploid cell (normal cell) can be affected by various copy number events (comprising various duplications, deletions, whole genome doubling (WGD), and their combinations). We use an HMM, a generative model with latent copy number states, to generate the observed read ratios—HMM, as in HMMcopy Shah et al. (2006), is a well-known model for copy number profiling. Hence, we generate 18 configurations of simulated single-cell clustering (CONF 1 to CONF 18 shown in Fig. 5), where each color-coded cluster corresponds to a unique CNV pattern, i.e., state transition pattern in the Markov chain. We use 150 cells and 200 bins, where the sequence is divided into relative chromosomes, i.e., the proportions of the chromosome lengths in the sequence are the same as those in a human genome. Relative chromosomes account for the independence of copy number variations between chromosomes. For the details of the simulated data, see Appendix A.8.

**Figure 5.**
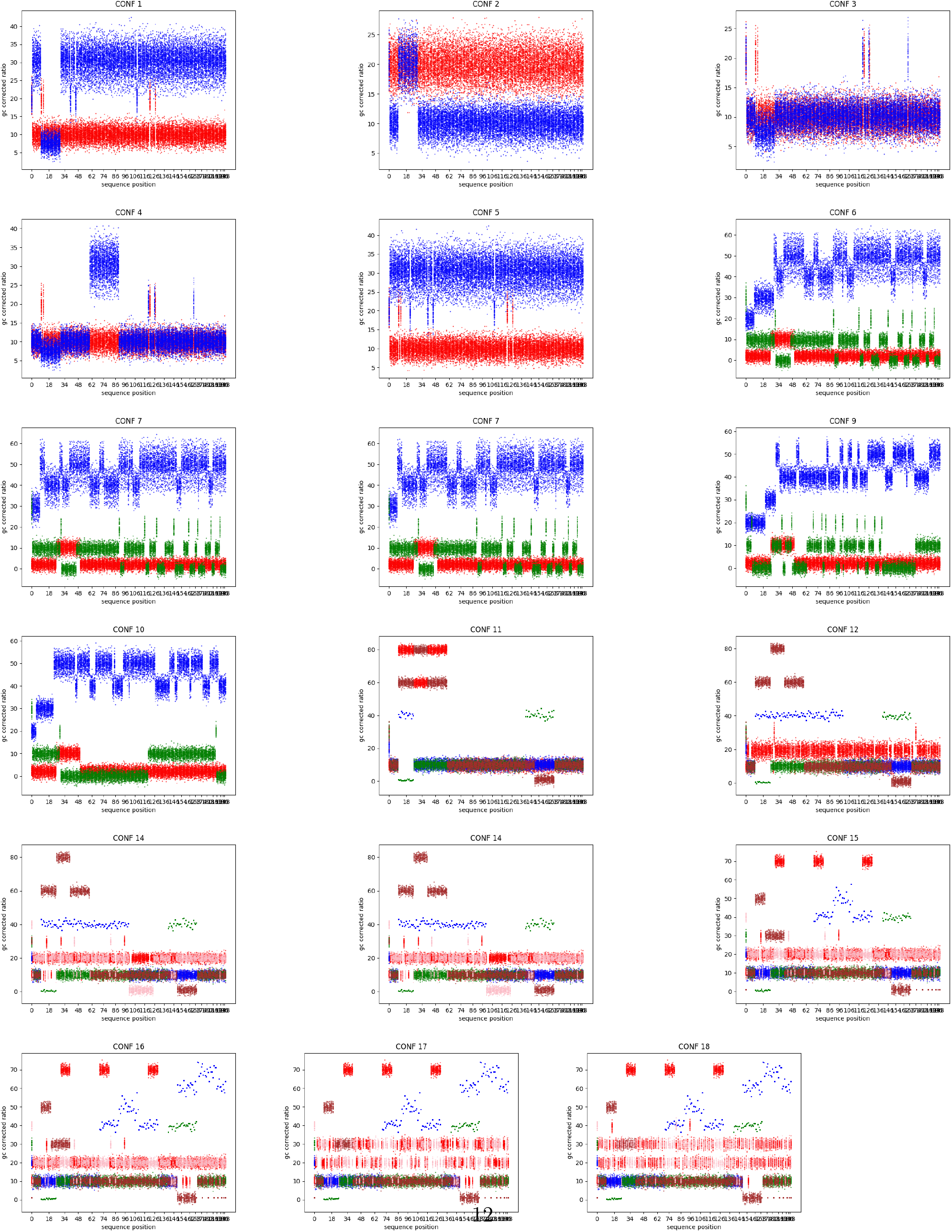
The datasets for CONF 1 to 18 are shown. CONF 1 to 5, CONF 6 to 10, CONF 11 to 13, and CONF 14 to 18 correspond to scenarios where 2, 3, 4, and 5 clusters are color-coded, respectively.

#### 3.1.2 Experimental Protocol

The VI framework allows for determining the number of clusters Bishop (2006); we run VI with a maximum number of clusters, and the framework results in zero probabilities for the extra clusters not detected by VI. We run VI using 50 different random initializations, as recommended by Blei et al. (2017). We choose the best VI run based on the highest ELBO.

For the Poisson version, we run VI for different numbers of clusters considering various random and informative initialization methods, e.g., k-means and Baum-Welch, as recommended by Blei et al. (2017). For each given number of clusters, we choose the best VI run based on the highest log-likelihood. Next, we select the number of clusters by an elbow selection method Johnson and Wichern (2007) based on the Deviance Information Criterion Spiegelhalter et al. (2002) over different numbers of clusters. We assess the clustering performance of CopyMix on 30 simulated datasets for each test case via V-measure Rosenberg and Hirschberg (2007). The V-measure, a value between 0 and 1, quantifies to which extent a cluster only contains data points that are members of a single class (the higher V-measure, the better clustering). Regarding inference of copy number profiles, we evaluate the performance of CopyMix on simulated data by calculating the proportion of discordant (unequal) position values between true and inferred vectors, as in de Souza et al. (2020), of copy number states. For biological data evaluation, V-measure is used to assess the clustering performance where the true clusters are assumed to correspond to the primary and metastatic cells; note that these may not be ground truth, but we believe there is certain concordance with the unknown ground truth. Regarding state inference, we compare the copy number profiles between the results from CopyMix and the original paper Leung et al. (2017).

For the Gaussian version, we compare the clustering results of CopyMix with the ground truth using V-measure for simulated data. For biological data evaluation, V-measure is used to compare our inferred clusters to the ones reported in the original paper Laks and McPherson (2019). In addition to merely comparing to the clustering result in the original paper, we examine if the report in the original article reveals meaningful clusters; we obtain this by calculating the CH Index Calinski and Harabasz (1974) and Silhouette Rousseeuw (1987) metrics on the raw data. Next, we use the likelihood ratio test to examine if the null hypothesis (CopyMix) hypothesis is rejected or not. Considering also SNV data, which are independent from CNVs, we run the SNV-CopyMix on the DLP data. Finally, we run GinkgoGarvin et al. (2015)—a state-of-art framework in copy number based clustering of single-cells by using circular binary segmentation method for copy number profiling—on the DLP data. For evaluating copy number profiles, we calculate the distance between the inferred and true copy number using total variation.

CH Index is the ratio of the sum of between-clusters dispersion and of inter-cluster dispersion for all clusters; the higher the score, the better the performance. Silhouette is calculated using two scores *a, b*, as shown below. Note that *a* is the mean distance between a sample and all other points in the same class. This score measures the closeness of points in the same cluster. And, *b* is the mean distance between a sample and all other points in the next nearest cluster. This score measures the distance of points of different clusters.

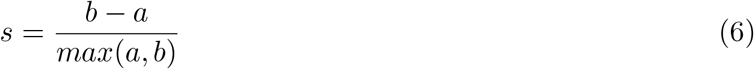

The *s* score is bounded between -1 for incorrect clustering and +1 for highly dense clustering. Scores around zero indicate overlapping clusters. The score is higher when clusters are dense and well separated, which relates to a standard concept of a cluster.

Finally, the likelihood ratio test assesses the goodness of fit of two competing statistical models based on the ratio of their likelihoods. In our experiments, we examine how statically significant the winning model is; by the winning model, we mean the one with the higher likelihood. The significance is measured by p-value; a high p-value indicates that the null hypothesis is preferred over the winning hypothesis.

### 3.2 Experimental Results for Poisson Version

We perform experiments on simulated data and biological data, and we evaluate a sequential version of CopyMix to verify the superiority of single-phase inference over the sequential framework. **Simulated data** Here, we evaluate CopyMix, and compare it to the initialization methods used by VI on 30 simulated datasets for each test scenario. The comparison is to illustrate, firstly, that VI results are not already achieved by the initialization methods, and secondly, that VI can recover the poor initialization performance and achieve optimal performance. We select the number of cells, sequence lengths, rate parameters, transition probabilities based on Leung et al. (2017). We assume equal amplification on the read counts across bins since the bin size is high enough (200kb as in Leung et al. (2017)), meaning that the effect of unequal amplification is negligible. Figures 6, 7 and 8 provide summaries of V-measures and proportions of discordant positions; note that the median values in the boxplots, highlighted in red for initialisation method and in blue for CopyMix, aim to facilitate the comparisons. The figures illustrate the performance comparison of CopyMix and the chosen initialization method, by CopyMix, for different settings: copy number transition matrices and number of clusters (Fig. 3); sequence length (Fig.4); and number of cells (Fig. 5). For all the tests, CopyMix results in V-measure of 1 except for one case (Fig. 6) where V-measure is in an interval of [.85, 1] with the median being 1; proportions of discordant positions are mostly 0 and otherwise are in the interval of [0, .2]. These results are optimal as the copy number profiling errors are near 0 and clustering near 1 (or 100%).

**Figure 6.**
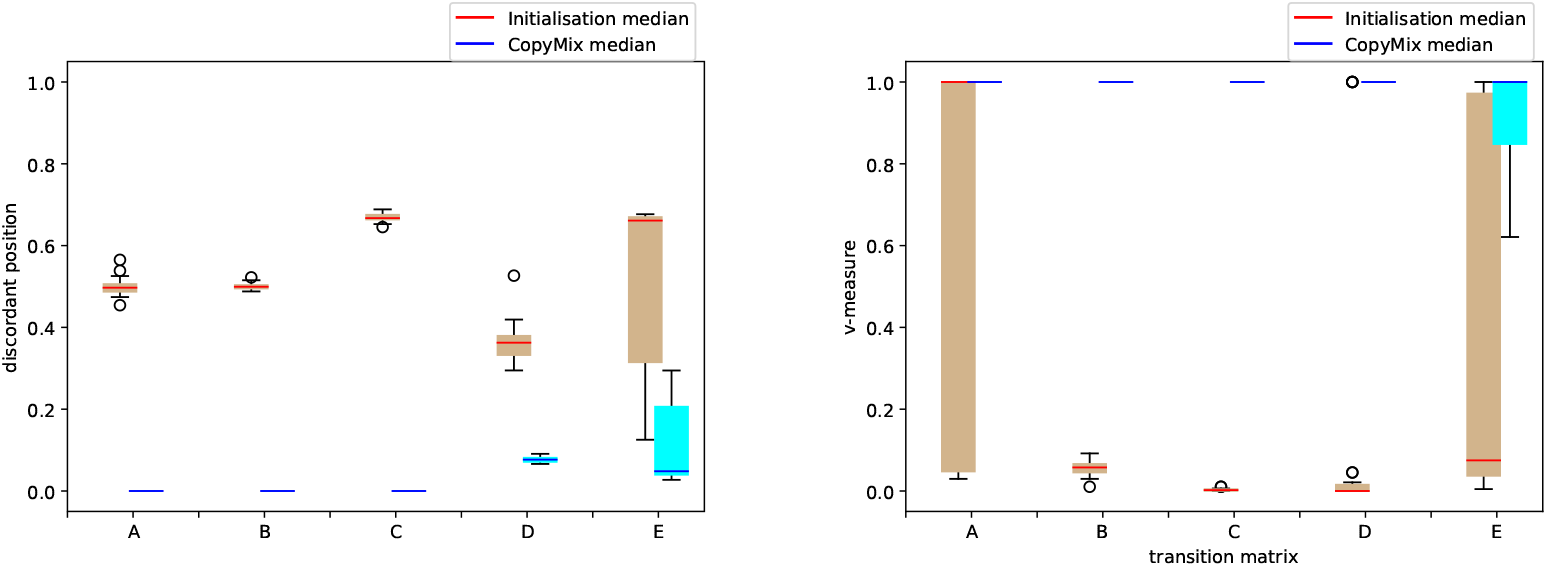
Performance of initialization methods (brown bars with red median values) and CopyMix (cyan bars with blue median values) across different sets of transition matrices. LHS: discordant positions between true and inferred vectors of copy number states. RHS: V-measure between true and inferred cell clustering assignments.

**Figure 7.**
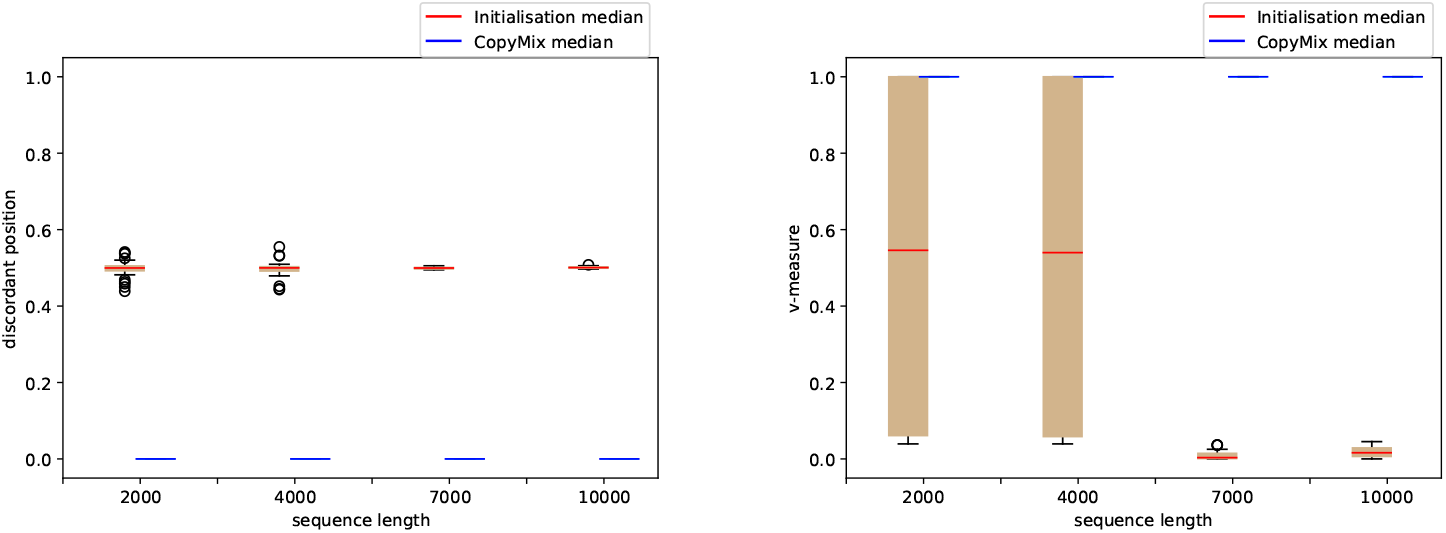
Performance of initialization methods and CopyMix, varying the sequence length. LHS and RHS figures, as well as color codes, are similar to Fig. 6.

**Figure 8.**
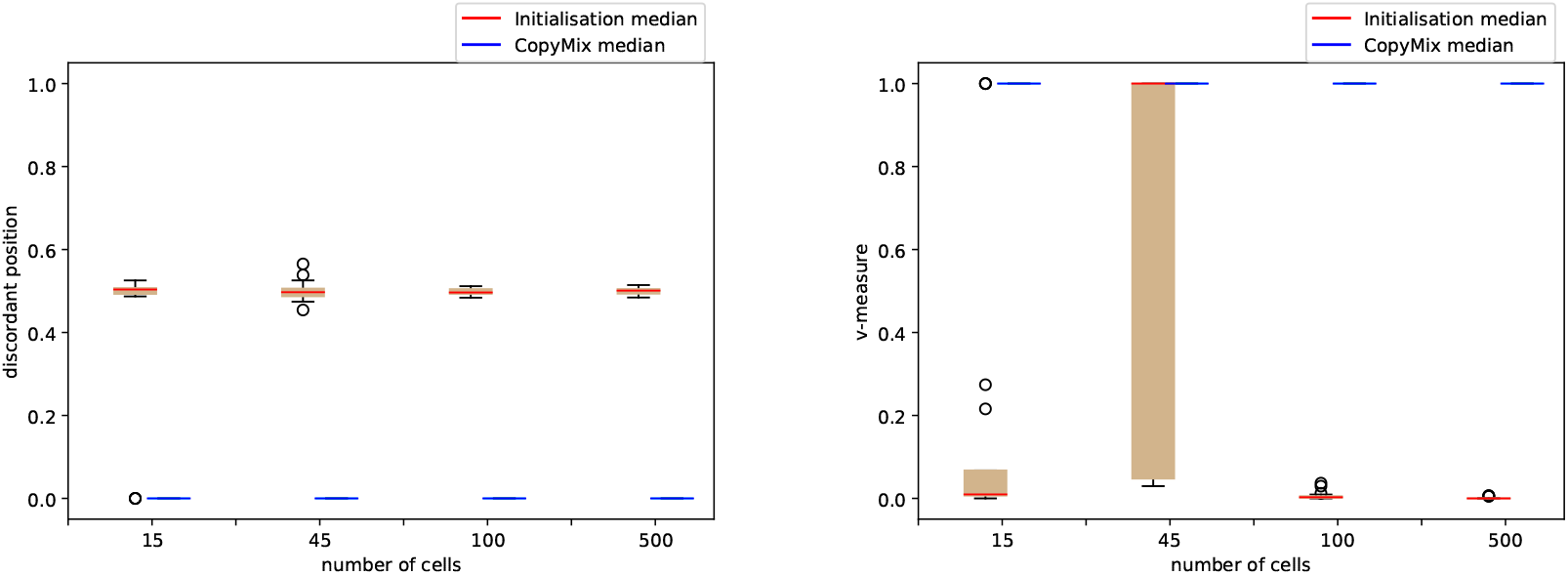
Performance of initialization methods and CopyMix varying the number of cells. LHS and RHS figures, as well as color codes, are similar to Fig. 6.

The results in Fig. 6 are obtained considering 45 cells—each with a sequence length of 800 bins and the mean value around 100—clustered into two and three clusters, wherein each clustering scenario, each cluster bears a different transition pattern than other clusters. The copy number transition patterns (forming the transition matrix of a Markov chain) vary across configurations A to E, which are orthogonal to each other; Configurations A to E are formed by combining the following state patterns.

- **single state**, a copy number state is inclined to stay at one certain state;
- **self-transition state**, a copy number state is inclined to stay at its current state for awhile;
- **oscillating state**, a copy number state changes shortly after being at a state;
- **spike state**, a sudden change of copy number state arises for a specific short range of positions in the copy number sequence;
- **scaled state**, a copy number changes by a multiplicative or additive constant.

The configurations A to E are: **A**, similar to chromosomes 11 in CRC1 and 4 in CRC2, (one cluster follows (**scaled state**); **B**, similar to chromosome 11 in CRC1 (one cluster follows **self-transition state** and the other one **single state**); **C**, similar to chromosome 3 in CRC1 (two clusters have identical transition matrices but one arrive at **spike state**); **D**, similar to chromosomes 18 to 21 in CRC2 (clusters combine **self-transition state, oscillating state**, and **scaled state**); **E**, two of the clusters have the transitions mentioned in case **C**, and, the third cluster has a combination of **oscillating state** and **single state**.

We can observe that, in Fig. 6, CopyMix has a perfect performance in terms of clustering and copy number inference for easier cases and also a high performance concerning the harder cases (**D** and **E**). Next, however not shown here, we investigate decreasing the rate parameter, *θ*_*n*_, for configuration **A** to see how our model handles shallow data as this can be a challenge in copy number profiling Zahn et al. (2017). Our model is robust towards lower rates up to *θ*_*n*_ = 3, without reducing the quality of performance concerning V-measure and discordant positions. As the third set of tests, we rerun the test using configuration **A**, but we increase the size of the sequences up to 10000 (inspired by the total length of the genomic bins from biological data in Leung et al. (2017)), see Fig. 7. We can observe that our model handles these sequences without any decrease in performance. Finally, Fig. 8 illustrates the perfect performance of CopyMix when varying the number of cells from 15 to 500 when, again, using configuration **A**.

#### Biological data

We evaluate CopyMix on the biological data from Leung et al. (2017). As aforementioned, we do not compare CopyMix to the earlier state-of-the-art methods due to their inclusion of ad-hoc processes in their framework; however, to evaluate the CopyMix results on the biological data, we use biologically verifiable results from the method in Leung et al. (2017). Using chromosomes 18 to 21, CopyMix clusters the cells into primary and metastatic groups, perfectly matching the corresponding primary and metastatic organs—V-measure is 1 w.r.t. the cells found in the two organs Leung et al. (2017). Adding chromosomes 3 and 8 to our test, metastatic tumor cells are clustered into two major subclusters as in Leung et al. (2017). As mentioned in Leung et al. (2017), amplifications on chromosomes 3 and 8, where some known genes are present, contribute to the detection of the metastatic subclusters. Note that, Leung et al. (2017) does not present a method to determine the threshold in hierarchical clustering, which results in the reported subclusters; moreover, there is no ground truth to verify these results.

Regarding the copy number profiling, Fig. 9 illustrates the copy number profiles inferred by CopyMix and the original work by Leung et al. (2017), here referred to as Navin. The heatmaps indicate a higher copy number state by a darker color. Note that in Navin, they perform ploidy scaling, which contributes to scaling differences compared to the results of CopyMix. Moreover, the Navin heatmap, here, illustrates an average of the heatmap in the original work across the cells since there is no single copy number profile per cluster. That is, Navin, unlike CopyMix, lacks a single copy number profile per cluster, revealing the copy number variation pattern behind each cluster. CopyMix detects the breakpoints (copy number changes) similar to Navin, with an accuracy of 76%, using the breakpoint margin (of a size corresponding to 5% of the sequence length). Again, there is no ground truth to verify the Navin results from Leung et al. (2017).

**Figure 9.**
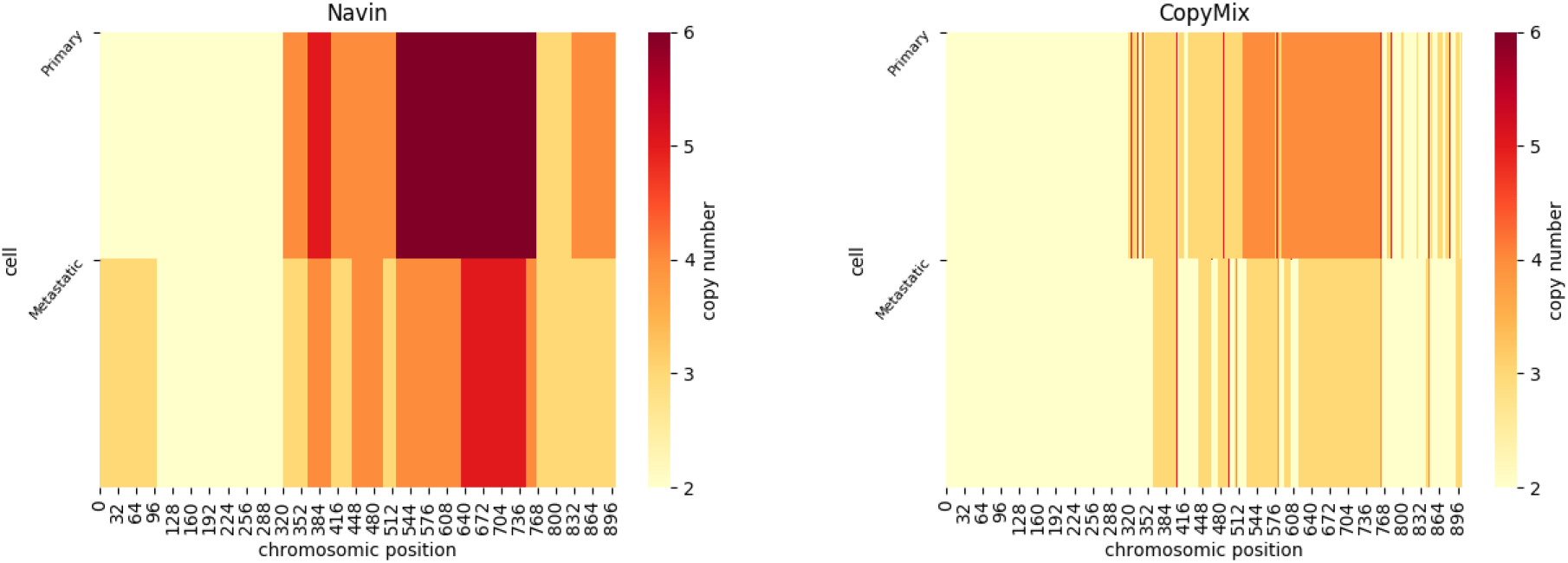
Copy number profiling comparison between the results from Navin Leung et al. (2017) (*LHS*) and CopyMix (*RHS*) for both primary and metastatic clusters.

#### A sequential CopyMix

Applying a sequential version of CopyMix (1. performing copy number profiling, using EM for HMMs; 2. performing clustering over the inferred copy numbers, using EM for a mixture model), resembling Shah et al. (2006); Vitak et al. (2017) but excluding their ad-hoc processes, on the biological data, results in poor clustering results (V-measure of .24) while, as mentioned, CopyMix gains V-measure of 1. This result confirms that simultaneous detection of copy number profiles and clustering is superior to the sequential approach. In particular, we conclude that methods that use EM for HMMs, using HMMcopy Shah et al. (2006) framework, benefit from the multiple ad-hoc transformations within the HMMcopy framework, much more than the pure computational model, e.g., HMM. Adding multiple ad-hoc rules within such computational models lacks elegance, integrity, and potential model-based approach benefits.

### 3.3 Experimental Results for Gaussian Version

We perform experiments on simulated data and biological data. For the biological data we evaluate CopyMix against the original paper, referring to as DLP, using CH Index, Silhouette, and likelihood ratio test.

#### Simulated data

We evaluated the performance of CopyMix on 16 simulated dataset configurations; see Fig. 5. In Table 1, we report V-measure for each configuration. We can observe that CopyMix, except for CONF 5, has a good clustering performance with V-measures ranging from 87% to 100%, Table 1. The exception, i.e., CONF 5, consists of two simulated genomes where one is a whole-genome duplication (WGD) of the other, *a whole-genome duplication pair*. Typically, when a genome undergoes a whole-genome duplication, the copy numbers across the sequence are scaled to a larger value, i.e., usually, diploid becomes tetraploid. This issue of CopyMix in distinguishing the ploidy can be explained partially by the notion of *unidentifiability*—inability to derive the true labels in unsupervised learning—of the transition matrix in a Markov chain Murphy (2012).

**Table 1:** V-measure (clustering result %) and total variation (copy number inference result 0.0) for CONF 1 to CONF 18.

|  |  |  |  |  |  |
| --- | --- | --- | --- | --- | --- |
| **CONF 1** | **CONF 2** | **CONF 3** | **CONF 4** | **CONF 5** | **CONF 6** |
| 100% — 0.09 | 100% — 0.05 | 100% — 0.05 | 100% — 0.20 | 0% — 0.11 | 100% — 2.45 |
| **CONF 7** | **CONF 8** | **CONF 9** | **CONF 10** | **CONF 11** | **CONF 12** |
| 100% — 2.45 | 100% — 2.45 | 100% — 2.36 | 100% — 2.45 | 87% — 1.02 | 100% — 0.96 |
| **CONF 13** | **CONF 14** | **CONF 15** | **CONF 16** | **CONF 17** | **CONF 18** |
| 100% — 0.93 | 88% — 1.20 | 87% — 0.96 | 100% — 1.03 | 100% — 1.04 | 100% — 1 |

Another important observation from Table 1 is that Copymix is robust w.r.t. increasing the number of clusters and the complexity of copy number patterns. We, however, notice a possible limitation of our purposeful modeling approach employed in CopyMix, i.e., each bin for each cell has a mean equal to the product of the cell-specific rate and the underlying latent copy number at that bin. Namely, despite providing strength in differentiating clusters, the multiplicative structure of copy number state and cell-specific rate can produce the same means for different clusters, e.g., 2 *×* 4 = 1 *×* 8, where 2 and 1 are the copy numbers of two clusters, and 4 and 8 are two cell rates corresponding to those two clusters, respectively; therefore, the higher cell rates may not be assigned to the higher copy number states because of this unidentifiability issue. The other source of clustering error is when the clusters have too overlapping sequence patterns, i.e., copy number profiles (CONF 14 and CONF 15 in Fig. 5). The cluster-overlapping in CONF 16 to CONF 18 is less than in CONF 15, hence perfect performance. As it may be hard to detect, we highlight that CONF 17 and CONF 18 differ in CNV in the last third part of the genome concerning clusters red and pink.

Finally, total variations between the true copy number and those inferred by CopyMix are shown in Table 1. CopyMix reveals small errors (distances to the ground truth copy numbers). However, we observe that slightly larger errors begin to manifest by increasing the number of clusters. Also, we notice that the error slightly increases when there are many breakpoints in the copy number profiles (the blue-colored genomes in CONF 6 to 10). This is because the possibility of differing copy numbers increases as copy number breakpoints increase.

#### Biological data

We evaluated CopyMix on the DLP data Laks and McPherson (2019), where each genome is partitioned into 6206 bins (bin size of 500K) ^3^. The clusters should agree with cell lines; no cluster contains cells from two different cell lines. It turns out that CopyMix succeeds in clustering the main clusters since the V-measure when comparing CopyMix clusters with the cell lines is 98%. The V-measure of overall CopyMix clusters based on those reported by the original clusters is 67%.

Using the LR test, the number of clusters, nine, reported by the original paper is statistically non-significant, i.e., CopyMix as a null hypothesis is favored over the method in the original paper. Assuming that each cluster follows an empirical Gaussian distribution—the mean and variance can be calculated by maximum likelihood estimate, nine clusters result in a log-likelihood of −8980284.42, and CopyMix log-likelihood is −9023905.94. The test results in *p*-value one, i.e., the original likelihood, is negligibly higher than that obtained by CopyMix.

Table 2 shows that CopyMix outperforms the original paper w.r.t. clustering according to CH Index and Silhouette metrics. We also evaluated Silhouette for fewer clusters reported by the original paper; these clusters are obtained according to the reported phylogenetic tree in the original paper Laks and McPherson (2019). Regarding Silhouette, even though the CopyMix score, 0.49, is not the perfect score, it is far better than those obtained by the method in the original paper. These metrics show that the method in the original paper has detected overlapping clusters, which is not sufficiently supported by the data. The SNV-CopyMix did not detect more clusters due to shallow data; for the details, see Appendix A.11. Note that the clusters reported by the original paper are not ground truth; hence, we cannot conclude that any diverging result is poor.

**Table 2:** CH Index and Silhouette for different numbers of clusters.

| Method | # of clusters | CH index |  |
| --- | --- | --- | --- |
| Original Paper | 9 | 462 |  |
| Cell lines from Original Paper | 3 | 623 |  |
| CopyMix | 4 | **1082** |  |
| Method | # of clusters | Silhouette | Description |
| Original Paper | 9 | 0.06 | Overlapping clusters |
| Original Paper w/o 2 minor clones | 7 | 0.21 |  |
| Original Paper w/o 4 minor clones | 5 | 0.25 |  |
| CopyMix | 4 | **0.49** | Separated clusters |

Finally, we compared CopyMix’s clustering results with those of Ginkgo. Unlike CopyMix, Ginkgo applies multiple ad-hoc processes, such as outlier removal from cells and bins and various corrections to the processed data. After removing 80% of the cells (717 out of 891 cells), Ginkgo detects three major clusters using hierarchical clustering (V-measure of 98% like CopyMix). CopyMix is more powerful than Ginkgo because CopyMix does not need to remove 80% of the data to obtain the V-measure of 98%. For more details, see Appendix A.9.

Regarding the copy number profiling, we obtained a total variation of 91 when comparing the copy number reported by CopyMix and the original paper. We can calculate the ratio of this total variation over the sequence length, 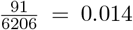, indicating a low error or relative total variation w.r.t. the sequence length (6206 bins of the genome). Since Ginkgo does not report the copy number profiles more than visually (see Appendix A.9), it is impossible to calculate its total variation.

## 4 Concluding remarks

We introduced CopyMix, a novel mixture model to jointly perform single-cell copy number profiling and clustering. CopyMix, while enjoying the advantages of Bayesian inference, addresses the following issues: (1) the sequential treatment of copy number profiling and single-cell clustering, prone to introduce clustering artifacts; (2) the labor of HMMcopy parameter tuning; (3) the lack of clonal copy number profiling by MHMMs; (4) the complicated posterior approximation when deriving VI for HMMs.

We evaluated our approach on both simulated and biological data, which indicated that CopyMix performs well when estimating both single-cell clustering and the corresponding copy number profiles.

CopyMix can be extended for phylogenetic inference, improving clustering, and augmenting and refining the model. Designing an allele-specific copy number based model can enrich the DLP results even more, and, finally, modeling biallelic deletions and aneuploidy, inspired by the recently proposed CopyKat Gao et al. (2021), is also a biologically desirable task.

## Author contributions statement

N.S. partially designed CopyMix, partially developed the mathematics, fully implemented CopyMix, fully wrote the manuscript, partially selected the evaluations metrics, and fully conducted experiments on CopyMix. C.P.E.S. partially designed CopyMix, partially developed the mathematics, helped with the design and evaluation of the study, partially wrote and reviewed the manuscript. A.R. provided useful suggestions for the evaluation of CopyMix and was involved in the model discussions. H.K. provided SNV emissions for the SNV-CopyMix and provided the main text for the section “The likelihood model for SNVs”. H.T. ran Ginkgo on the biological DLP data. J.L. fully supervised and reviewed the paper, was involved in the entire project, and partially wrote the manuscript.

## A Appendix Derivation details

### A.1 Step 2 of Section 2.1

In this section, we provide the calculation of each decomposed term of log *P*(**Y, Z, C, Ψ**). Each term of log *P*(**Y, Z, C, Ψ**) presented in Step 2 of Section 2 is calculated as follows: 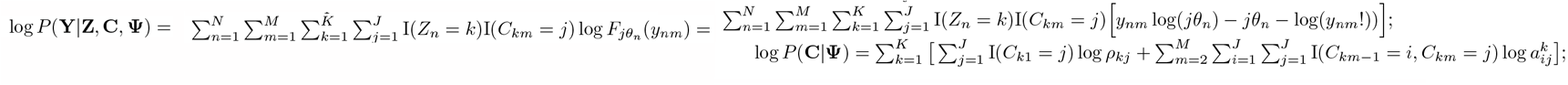

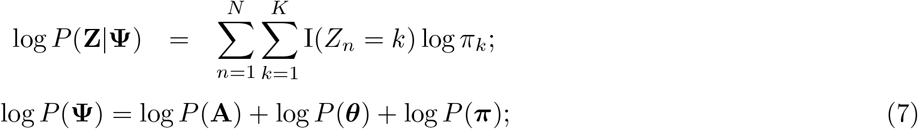

where,

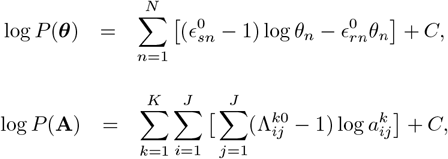

and

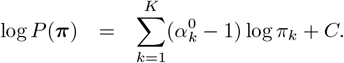

### A.2 Step 3 and 4 of Section 2.1

In this section we first present the details of the step 3 of section 2, i.e., the derivation of each posterior update equation (VD computation). We then summarise them in the form of a pseudo code, covering the details of step 4 in Section 2. For convenience, we use +≈ to denote equality up to a constant additive factor.

*Update equation for* ***π***: 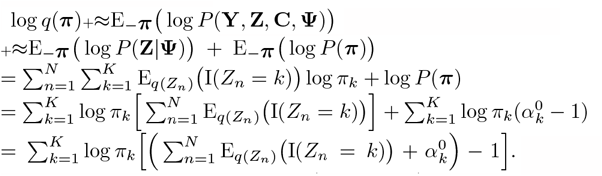.Let I(·) be the indicator function. *q*(***π***) is a Dirichlet distribution with parameters *α* = (*α*_1_, …, *α*_*K*_), where

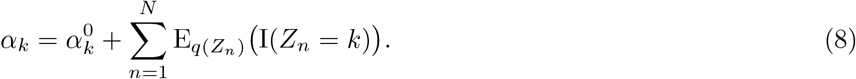

*Update equation for Z*_*n*_:

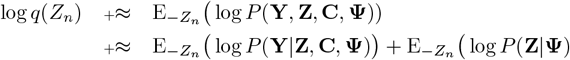

Let *y*_*nm*_ be the observed number of reads aligned to bin *m* for cell *n* corresponding to the random variable *Y*_*nm*_ and let *D*_*n,m,j*_ ≡ *y*_*nm*_ log *θ*_*n*_ + *y*_*nm*_ log *j* − *jθ*_*n*_ − log(*y*_*nm*_!). Note that log *P*(**Y**|**Z, C, Ψ**) and log *P*(**Z**|**Ψ**) can be written as the sum of two terms, one that depends on *Z*_*n*_ and one that does not, i.e., log 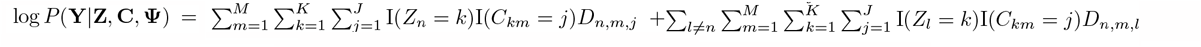 and

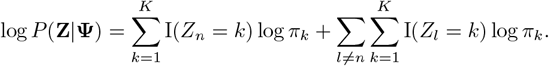

Consequently,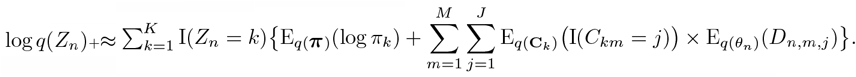

The distribution *q*(*Z*_*n*_) is a Categorical distribution with parameters ***π***_*n*_ = (*π*_*n*1_, …, *π*_*nK*_), where

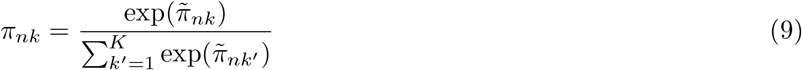

With

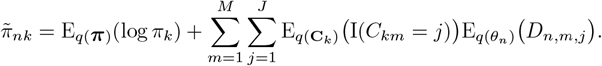

*Update equation for θ*_*n*_:

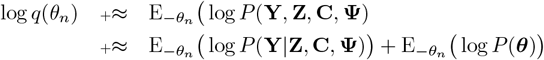

We can write log *P*(**Y**|**Z, C, Ψ**) as in Eq. A.7 and log *P*(***θ***) which depends on *θ*_*n*_ as,

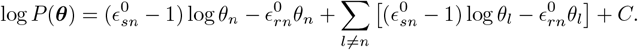

There fore, 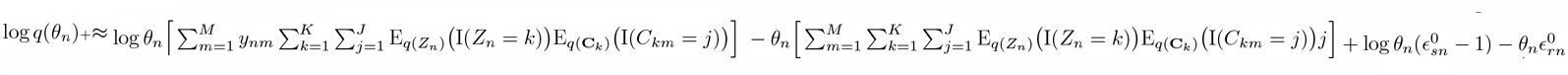.

Note that 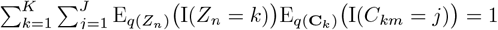.

The distribution *q*(*θ*_*n*_) is a Gamma distribution with shape and rate parameters:

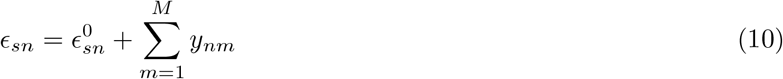

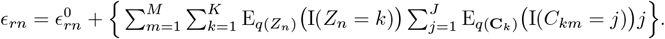

*Update equation for* 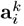:

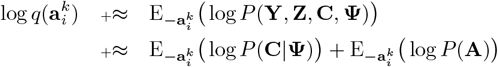

Disregarding the terms that do not depend on 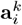 in log *P*(**C**|**Ψ**) and log *P*(**A**), we obtain: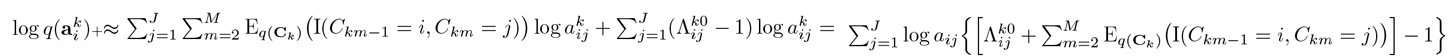

The distribution 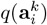 is Dirichlet with parameters 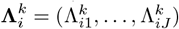, where

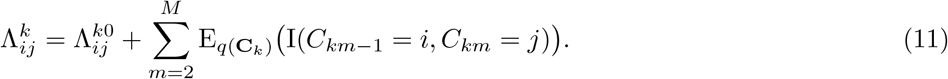

#### Update equation for **C**_k_

In the calculation of *q*(**C**_*k*_), the resulting distribution resembles the posterior probability of the hidden variables in an HMM and only differs in the normalization constant. Similarly to the approach adopted in MacKay (1997), we use a slightly different implementation of forward-backward algorithm. More specifically, we define a graph (one per cluster *k*) *G* = {*V, E*} with vertices *V* = {*C*_*mj*_ : *m* ∈ [*M*], *j* ∈ [*J*]}, having weights denoted as *w*_*k*_(*C*_*mj*_), and edges *E* = {*C*_*m*−1*i*_*C*_*mj*_ : *m* ∈ {2, …, *M* }, *j* ∈ [*J*]}, having weights denoted as *w*_*k*_(*C*_*m*−1*i*_*C*_*mj*_). Note that for the simplicity of notation we use *C*_*mj*_ instead of *C*_*km*_ = *j* for edges and vertices. We define forward 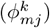 and backward 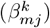 quantities similar to HMMs Bishop (2006). Instead of using the terminologies of transition and emission probabilities, we formulate a corresponding algorithm in terms of the weights of the graph, *w*_*k*_(*C*_*m*−1*i*_*C*_*mj*_) and *w*_*k*_(*C*_*mj*_). The initial transition probability can be defined as *w*_*k*_(*C*_0_*C*_1*j*_). Notice that *k* indicates the cluster which the graph belongs to. We calculate forward and backward quantities using the weights, and there is no need to normalize the graph weights since a normalization is performed later when calculating the VD. Having the weights, we compute the two posterior probabilities *q*(*C*_*km*_ = *j*) and *q*(*C*_*km*−1_ = *i, C*_*km*_ = *j*); note that they are normalized by summing over all *j* ∈ [*J*]. McGrory and Titterington (2009a) derive the exact normalization of the graph weights, for a similar but less complex graphical model; however, this is unnecessary since the normalization can be performed later, as we do.

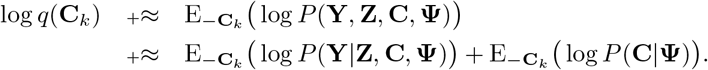

We can write log *P*(**Y**|**Z, C, Ψ**) as the sum of two terms, one that depends on **C**_*k*_ and another that does not, that is, 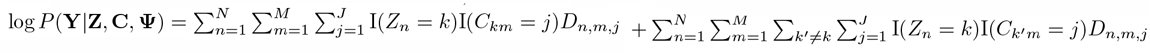.

Let *f*(*k*) be a function of *k* as 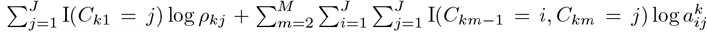, We can write 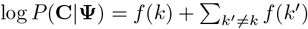.

Thus, disregarding the terms that do not depend on **C**_*k*_ we obtain: 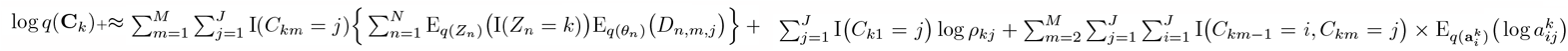. Calculations regarding di-rected graph are as follows: 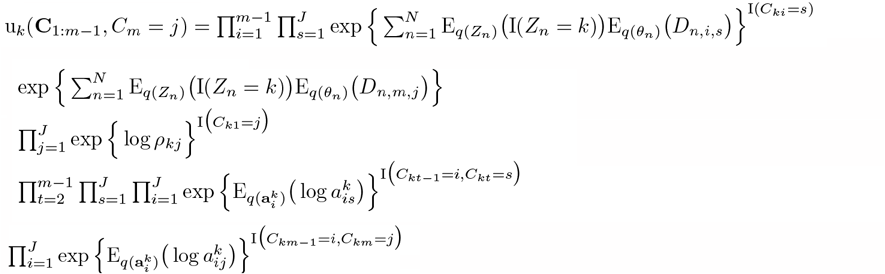 and

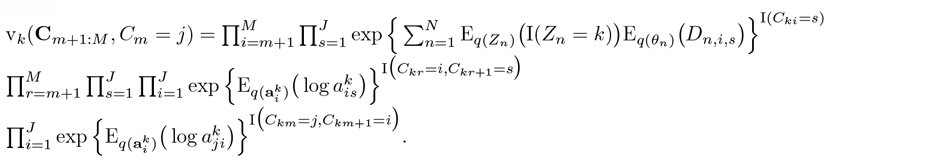

We define 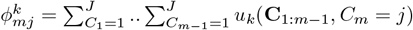 and 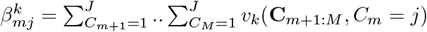

The graph weights are (we assume *w*_*k*_(*C*_0_*C*_1*j*_) to be fixed):

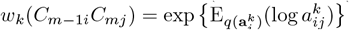

and

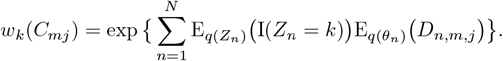

Note that we skip writing *i, j* in the calculations to make them short and more readable:

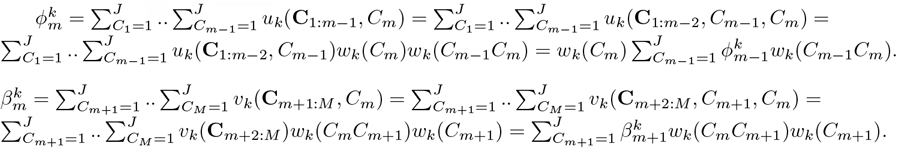

We now calculate the posteriors: 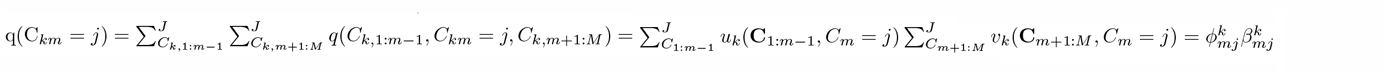

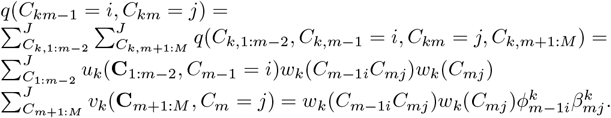

Finally, the following are the resulted update equations. Subsequently, the detailed algorithm used in CopyMix is shown in Algorithm 2.

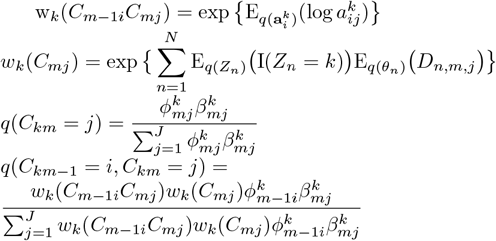

##### Algorithm 2 CopyMix: a Variational Inference Algorithm

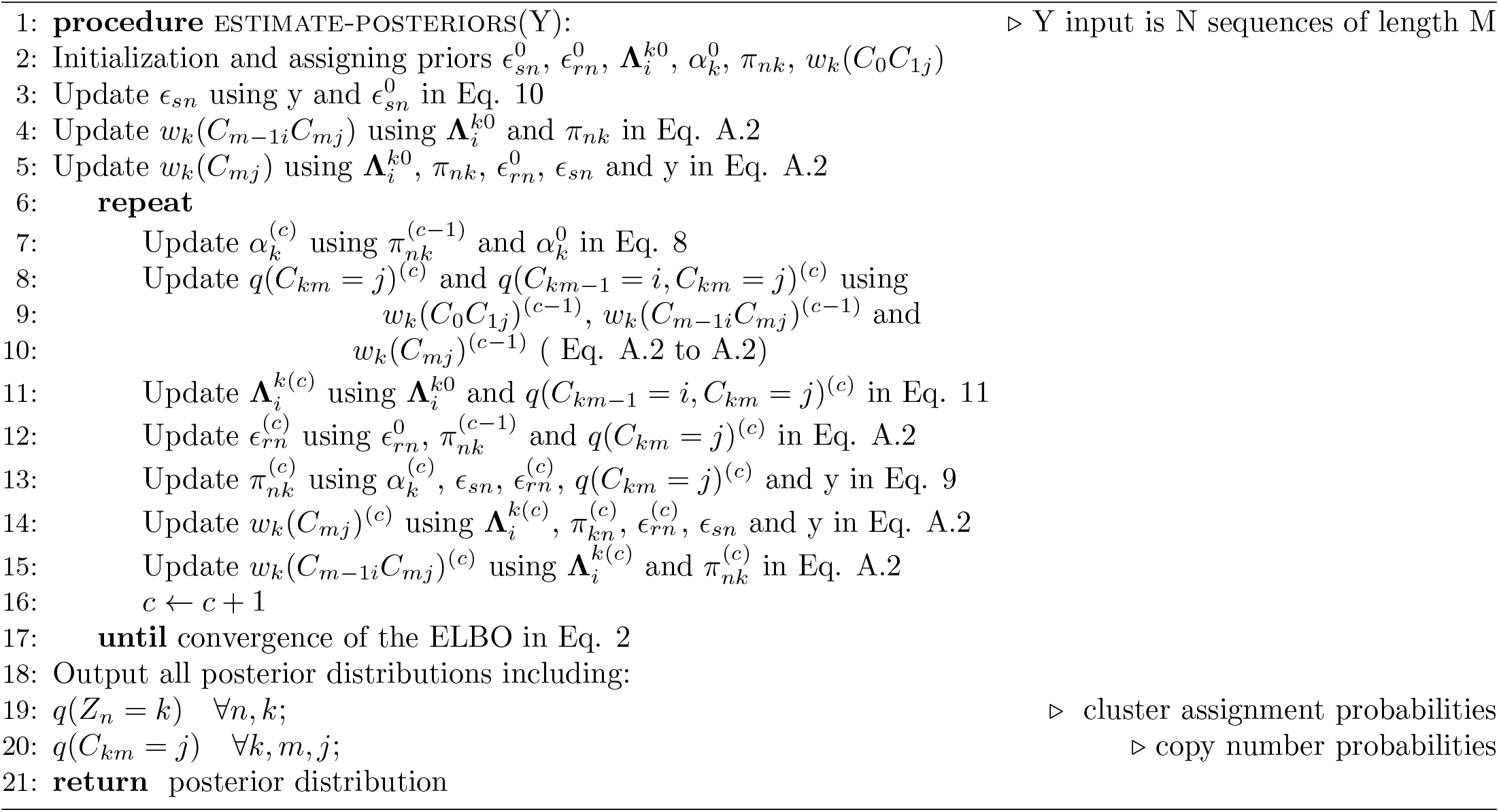

#### A.3 Calculating expectations for section 2.1

In this section we list the calculations of the expected values that are used in the update equations in the previous Appendix. Let **Φ** be the digamma function defined as 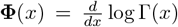 which can be easily calculated via numerical approximation. The values of the expectations taken with respect to the approximated distributions are given as follows where **Φ** is used for some of them.

- 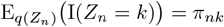
- Due to computational issues in calculating 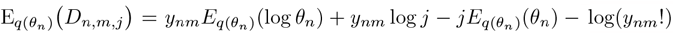, we can instead compute it using the fact that 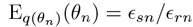 and 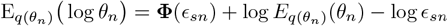.Therefore, we obtain 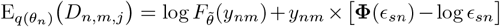, with 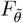 being a Poisson p.m.f. with rate parameter 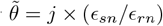.
- 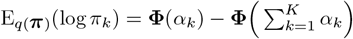
- 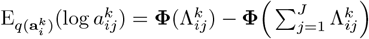
- 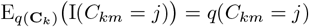 and 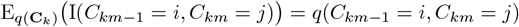, which are calculated in Eq. A.2 and A.2.

#### A.4 Step 2 of Section 2.2

The logarithm of the joint distribution of **Y, Z, C** and **Ψ** satisfies

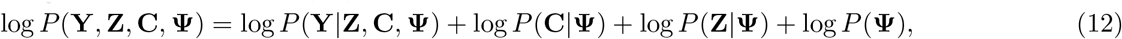

where

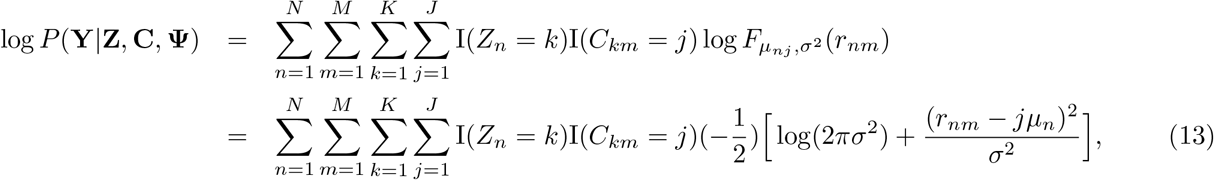

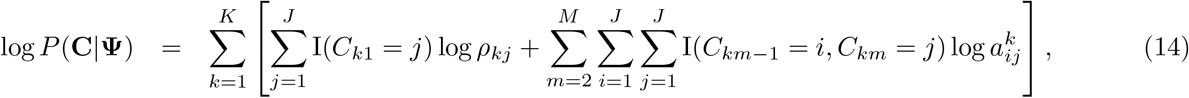

and

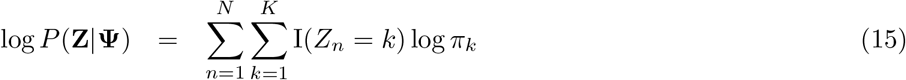

Moreover,

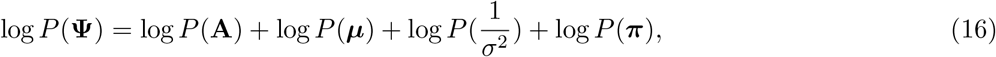

where, if B is the multivariate Beta function, i.e.,

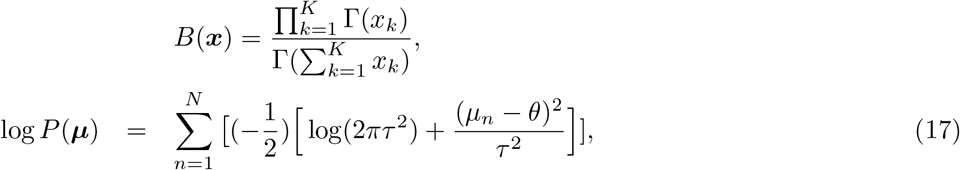

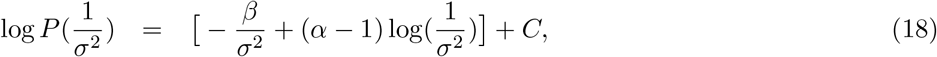

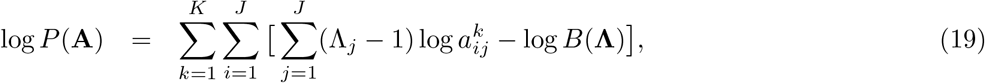

and

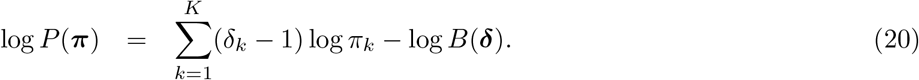

### A.5 Step 3 and 4 of Section 2.2

#### Update equation for π

Since (15) and (20) are the only terms in (12) that depend on ***π***, the update equation q^*\**^(***π***) can be derived as follows.

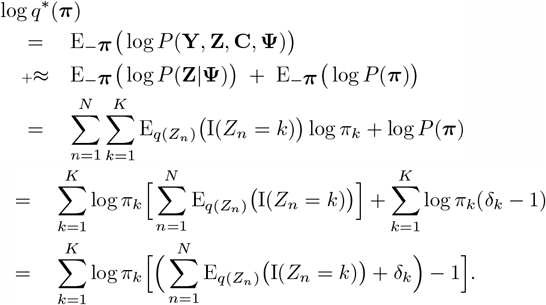

Therefore, q^*\**^(***π***) is a Dirichlet distribution with parameters 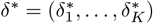, where

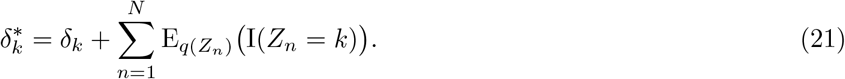

#### Update equation for Z_*n*_

Since (A.7) and (15) are the only terms in (12) that depend on Z_*n*_, q^*\**^(Z_*n*_) can be obtained as follows.

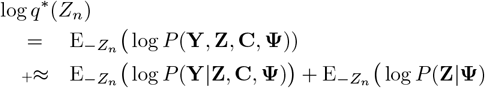

Note that log P(**Y**| **Z, C, Ψ**) and log P(**Z**| **Ψ**) can be written as the sum of two terms, one that depends on Z_*n*_ and one that does not, i.e.,

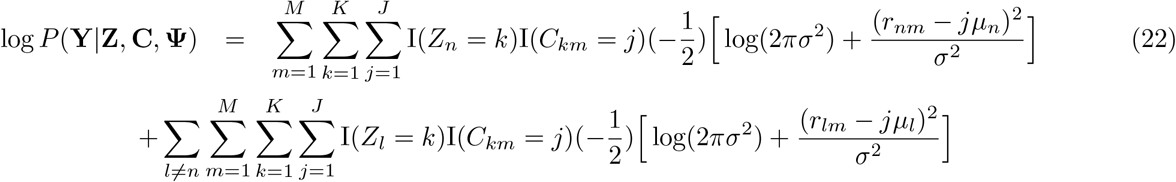

and

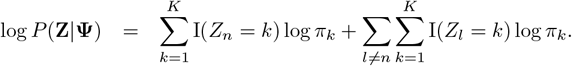

Consequently,

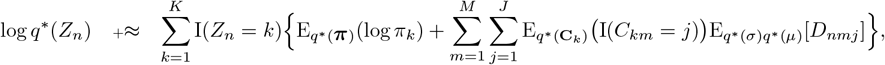

Where:

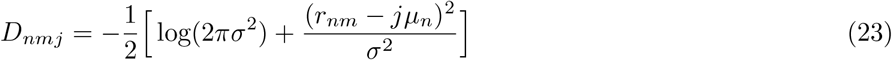

We conclude that q^*\**^(Z_*n*_) *∼* Categorical 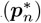 with parameters 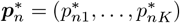 where

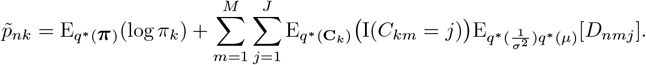

and

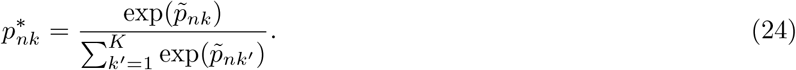

#### Update equation for μ

Since (A.7) and (17) are the only terms in (12) that depend on μ_*n*_, q^*\**^(μ_*n*_) can be obtained as follows.

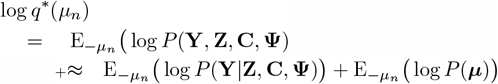

We can write log P(**Y**|**Z, C, Ψ**) as in Eq. A.7, and 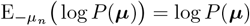, which is:

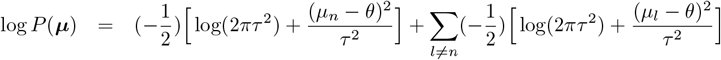

Therefore, log 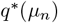.

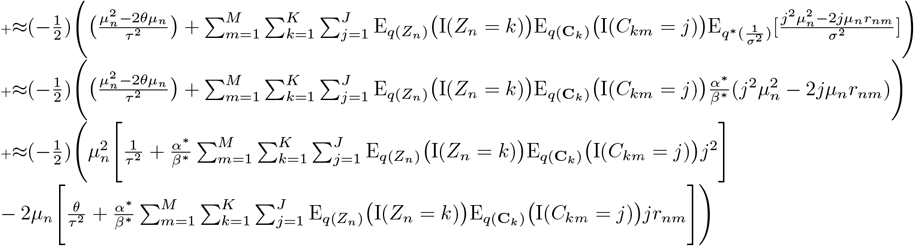

Thus, q^*\**^(μ_*n*_) is a Normal distribution with parameters:

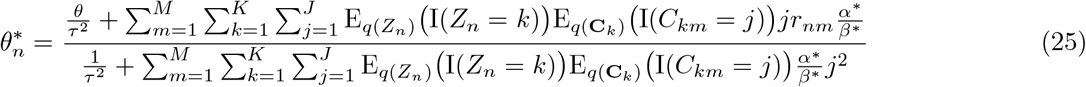

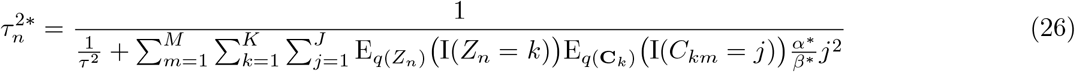

#### Update equation for 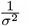

Similar to the previous calculations, we do as follows.

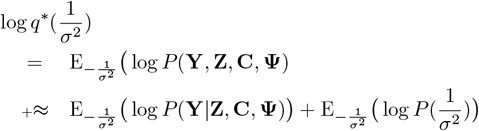

We can write log P(**Y Z, C, Ψ**) and log P(σ) as they are in Eq. A.7, 18. Putting them together, we achieve the following.

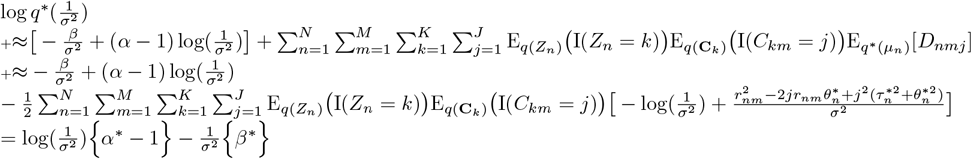

Therefore, 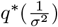 is a Gamma distribution with parameters:

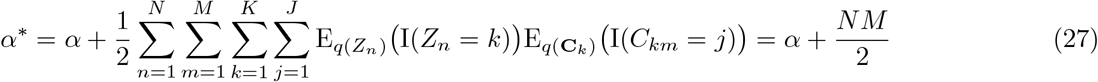

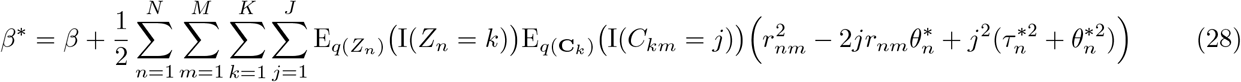

#### Update equation for 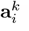

Because (14) and (19) are the only terms in (12) that depend on 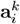, we calculate 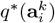 as follows.

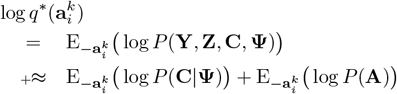

Disregarding the terms that do not depend on 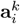 in log *P*(**C**|**Ψ**) and log *P*(**A**), we obtain:

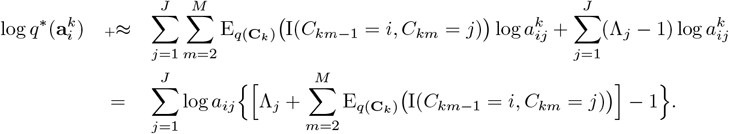

Therefore, 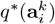 is Dirichlet with parameters 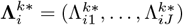 where

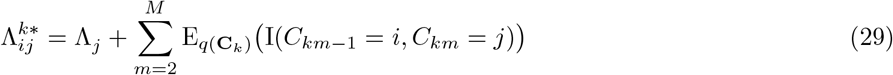

#### Update equation for C_*k*_

Since only (A.7) and (14) depend on **C**_*k*_, log q^*\**^(**C**_*k*_) can be calculated as follows.

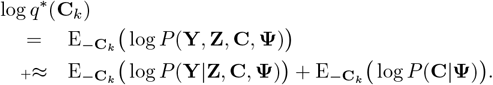

Similarly we can write log P(**C**|**Ψ**) as

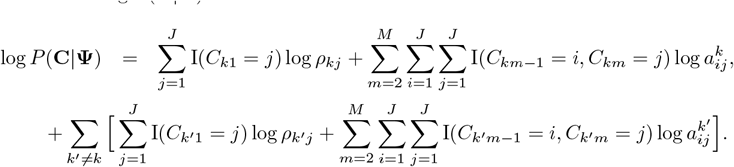

Thus, disregarding the terms that do not depend on **C**_*k*_ we obtain:

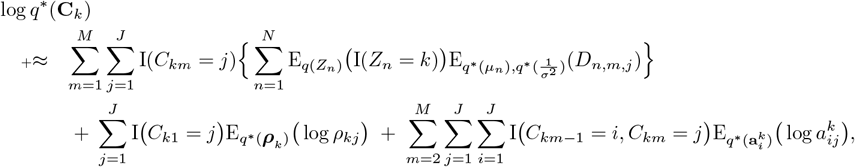

#### Calculations regarding directed graph

We define graph *G* = {*V, E*} where we have each vertex as *V* = {*C*_*mj*_ : *m* ∈ *M, j* ∈ *J*} with weight w(C_*mj*_) and each edge as *E* = {*C*_*mj*_*C*_*m*+1*j*_ : *m ∈ M, j ∈ J*} with weight *w*(*C*_*mj*_*C*_*m*+1*j*_). We define quantities of forward, *ϕ*_*mj*_, and backward, *β*_*mj*_, similar to HMM as below (the calculations are based on log *q*^*\**^(**C**_*k*_)):

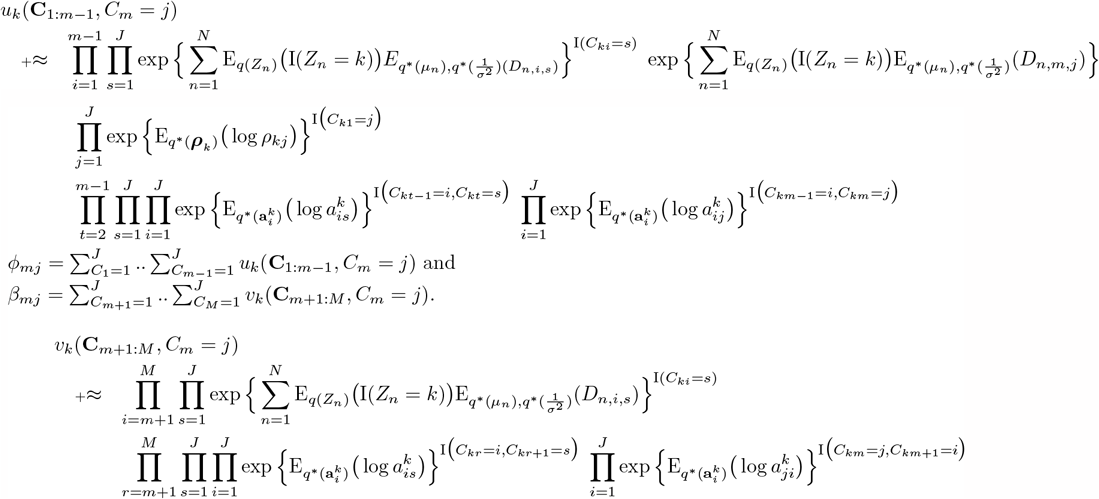

Instead of using terminologies of transition and emission probabilities we formulate those as weights of the graph; *w*_*k*_(*C*_*mj*_*C*_*m*+1*j*_) and *w*_*k*_(*C*_*mj*_) respectively. The initial transition probability can be define as *w*_*k*_(*C*_0_*C*_1*j*_) assuming the source starts at 1. Notice that *k* shows the cluster which the graph belongs to. We can calculate forward and backwards using dynamic programming; having those values we can compute the two posterior probabilities of 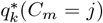 and 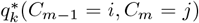 which are the expectations of the indicator functions; we then normalized them by summing over all *j ∈ J*. The graph weights are: 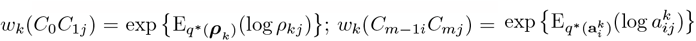 and 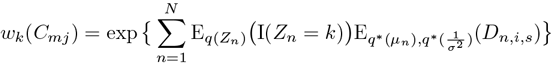.

Note that we skip writing *i, j, k* in the calculations to make them short and more readable:

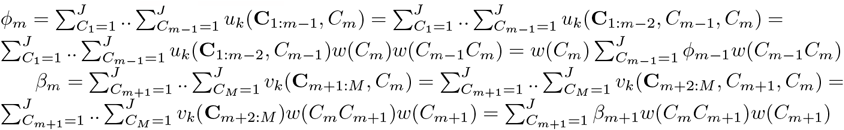

We now calculate the posteriors:

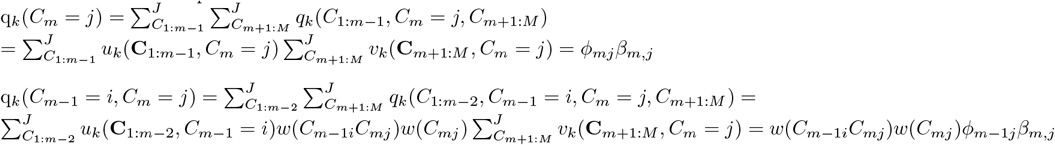

##### A.6 Calculating expectations and ELBO for Section 2.2

Let **Ψ** be the digamma function defined as

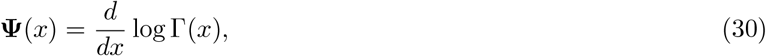

which can be easily calculated via numerical approximation. The values of the expectations above taken with respect to the approximated distributions are given as follows.

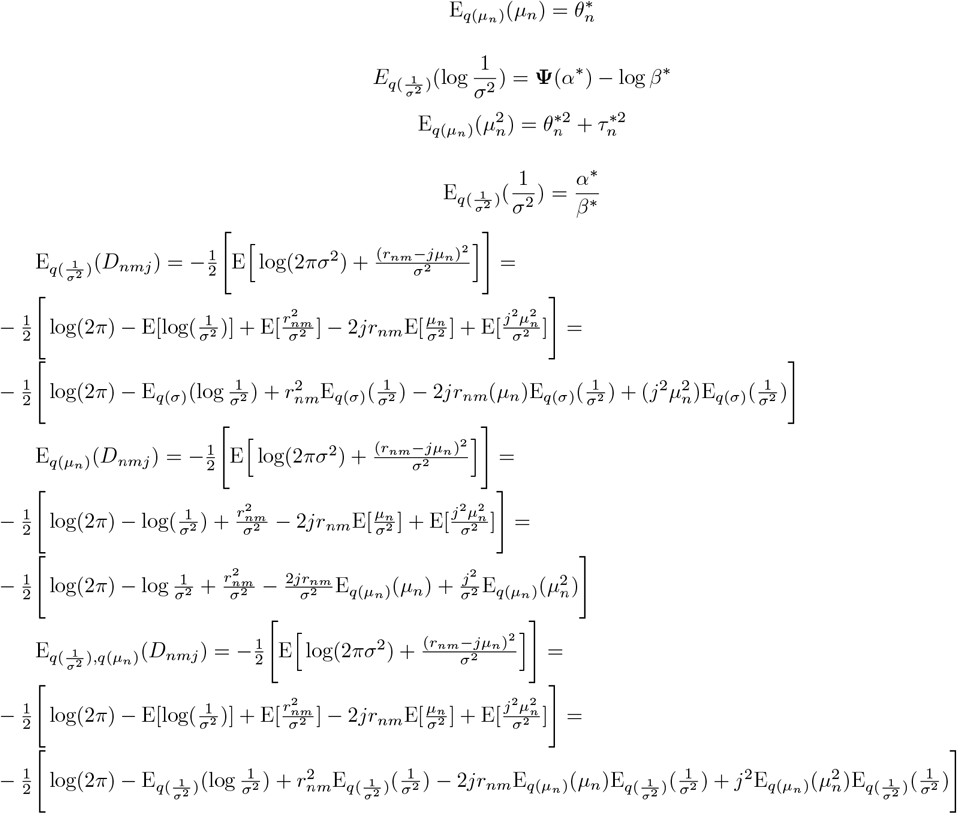

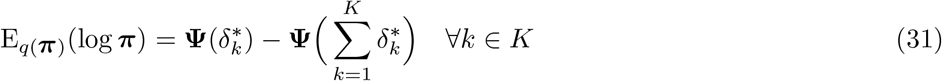

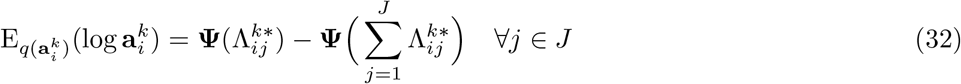

Using the results above regarding the expectations, we update the parameters of the approximated distributions iteratively.

We then conduct many iterations until the convergence of the ELBO in Eq. 2 which is calculated as below. An assumption in the ELBO calculation is that we ignore all the constants contributing in the ELBO value.

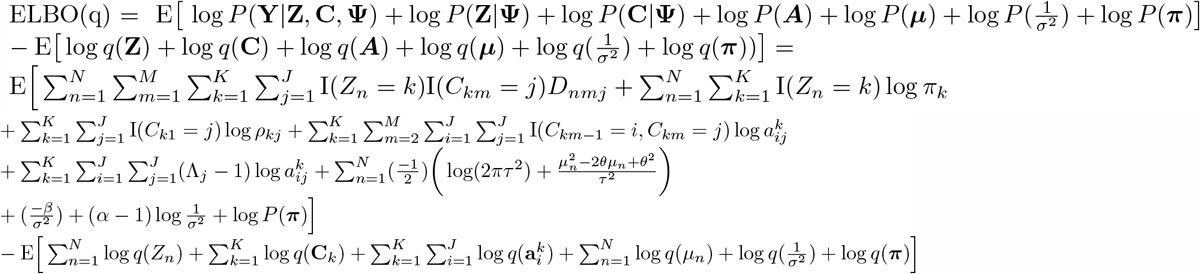

We calculate the E[log *q*(**C**_*k*_)] by decomposing it into initial and transition components. Note that the initial probabilities, *ρ*_*kj*_s, are fixed and, therefore, the term corresponding to that cancels out the corresponding term in E[log *P*(**C**|**Ψ**)]. The remaining term, the transition component, is calculated by the following using 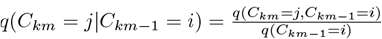, as below:

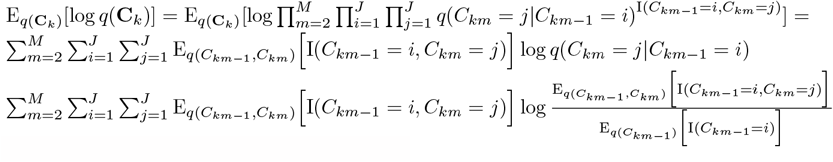

We calculate the E[log *p*(*Z*_*n*_|Ψ)] as below:

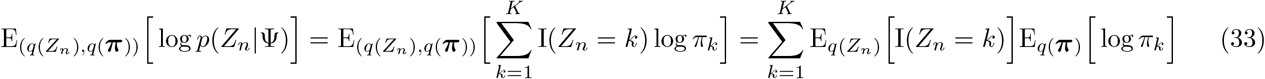

We calculate the E[log *q*(*Z*_*n*_)] as below. Note that the expectation of *q*(*Z*_*n*_) w.r.t. *q*(*Z*_*n*_) is equal to *q*(*Z*_*n*_). Also, we know that 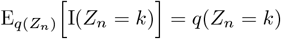.

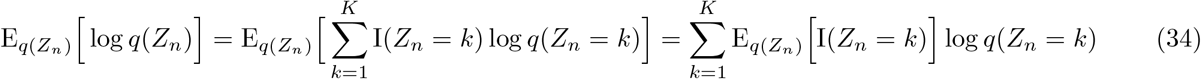

We calculate the E[log *P*(*μ*_*n*_)] as below:

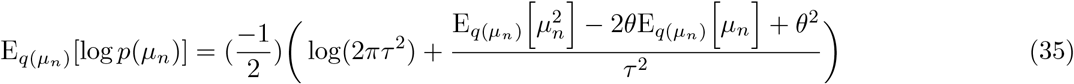

We calculate the E[log *q*(*μ*_*n*_)] as below:

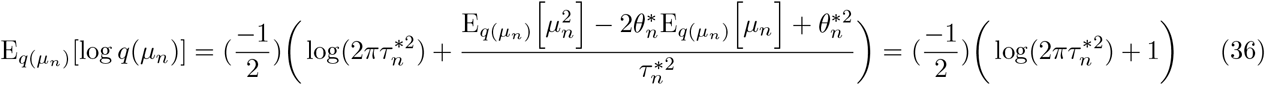

We calculate the E[log *P*(***π***)] as below:

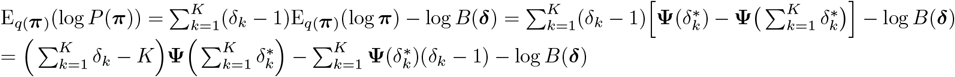

We calculate the E(log *q*(***π***)) as below:

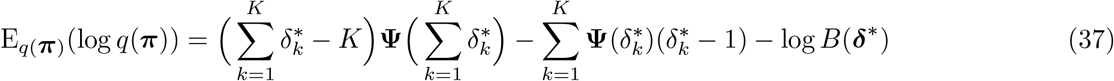

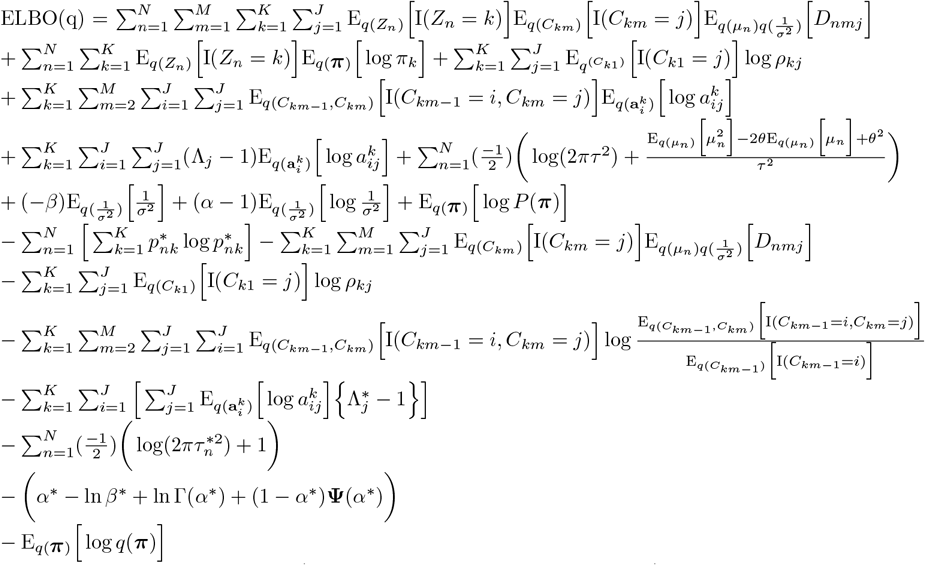

We now approximate the term,(*α*^*\**^ *−* ln *β*^*\**^ + ln Γ(*α*^*\**^) + (1 *− α*^*\**^)**Ψ**(*α*^*\**^)) in the above, by using approximations:

1. Stirling’s: ln Γ(*α*^*\**^) = (*α*^*\**^ *−* 1) ln(*α*^*\**^ *−* 1) *− α*^*\**^ + 1
2. digamma-approximation: 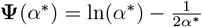
3. Also: ln 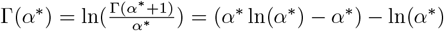

Putting 3 and 2 in the original formula we obtain:

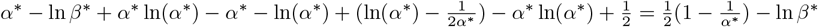

##### A.7 VI for SNV-CopyMix

Let **Ψ** be the set containing all the model parameters, i.e., **Ψ** = {**A, *μ***, *σ*, ***π***, *ξ}*, where

- **A** = {**A**_*k*_ for *k* = 1, …, *K}* with 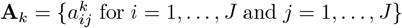 and *j* = 1, …, *J}*;
- ***μ*** = {*μ*_*n*_ for *n* = 1, …, *N }*;
- *σ*;
- ***π*** = (*π*_1_, …, *π*_*K*_);
- *ξ*.

In order to infer **Ψ** and the hidden states **Z** = (*Z*_1_, …, *Z*_*n*_), **C** = {**C**_1_, …, **C**_*K*_ }, and **S** = {*S*_1_, …, *S*_*K*_ }we apply the Variational Inference (VI) methodology; that is, we derive an algorithm that, for given data, approximates the posterior distribution by finding the Variational Distribution (VD),

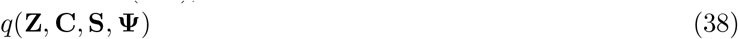

with smallest Kullback-Leibler divergence to the posterior distribution

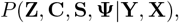

which is equivalent to maximizing the evidence lower bound (ELBO) given by

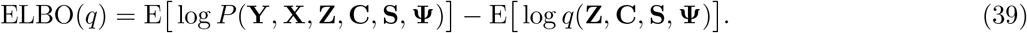

We consider the following prior distributions for the parameters in **Ψ**.

- 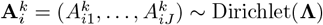
- 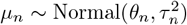. The conjugate prior concerning the mean of Normal distribution is Normal distribution.
- 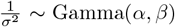. The conjugate prior concerning the precision of Normal distribution is Gamma distribution.
- ***π*** *∼* Dirichlet(***δ***).
- ξ_*k*_ *∼* Beta(*γ*_*k*_, *η*_*k*_).

In what follows we describe the main steps of the VI algorithm for inferring **Z, C, S** and **Ψ**.

#### Step 1. VD factorization

We assume the following factorization of the VD:

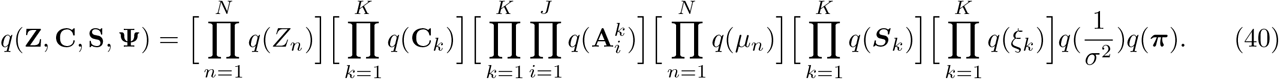

#### Step 2. Joint distribution of observed data, hidden variables, and parameters

The logarithm of the joint distribution of **Y, Z, C** and **Ψ** satisfies log *P*(**Y, X, Z, C, S, Ψ**) = log *P*(**Y**|**Z, C, Ψ**) + log *P*(**X**|**Z, S, Ψ**) + log *P*(**C**|**Ψ**) + log *P*(**Z**|**Ψ**) + log *P*(**S**|**Ψ**) + log *P*(**Ψ**),

where 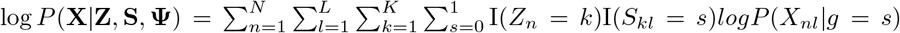We assume we have these probabilities given the genotype g being either mutation or non mutation.

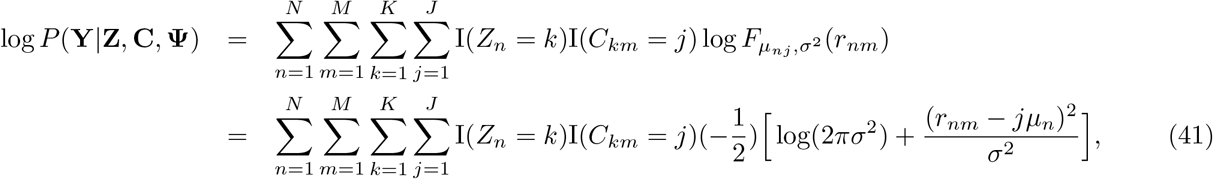

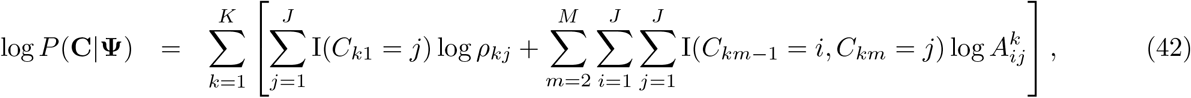

Note that ρ_*kj*_ is the initial probability in the MC which we fix.

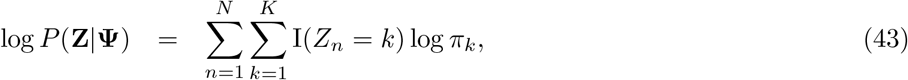

and

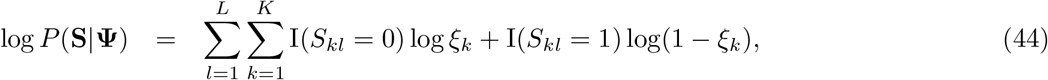

Moreover,

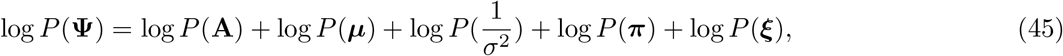

Defining, ℬ the multivariate Beta function, i.e., 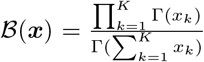, and 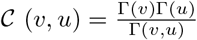, we have:

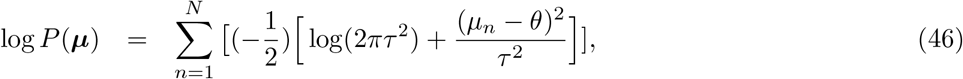

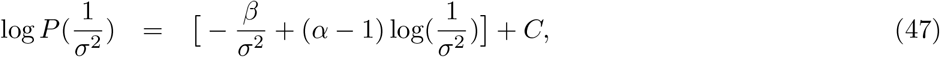

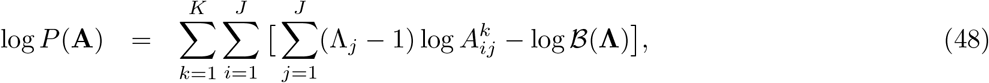

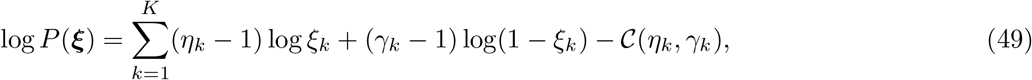

and

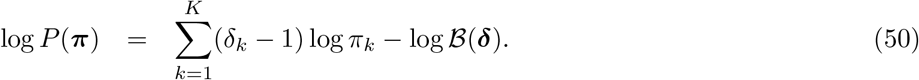

#### Step 3. VD computation by coordinate ascent

We now derive a coordinate ascent algorithm for the VD. That is, we derive an update equation for each term in the factorization (in step 1) by calculating the expectation of log *P*(**Y, X, Z, C, S, Ψ**) over the VD of all random variables except the one of interest. Below, the update equation is derived for each random variable. For convenience, we use +≈ to denote equality up to a constant additive factor. Note that **X** can have missing values at certain sites. Therefore, including this into the model, results in splitting **X** into **X**_**miss**_ and **X**_**obs**_. Moreover, we would need a Bernoulli variable, *ω*, which shows which of the two group is the case in the joint distribution. The joint distribution then takes the following form.

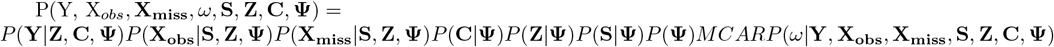

In the above, MCAR is referring to missing completely at random which is our assumption here, hence its probability is independent from all variational parameters. This means that this probability is a constant w.r.t. the variational parameters. We assume that *P*(**X**_**miss**_| *ω*, **S, Z, Ψ**) is constant and, therefore, consider *X* = *X*_*o*_*bs* in our applications. However, for the ease of notation, in our derivations we assume *X* is complete. Namely, we can, for the sake of readability, simplify the joint distribution as *P*(**Y, X, S, Z, C, Ψ**), where **X** refers to **X**_**obs**_.

**Update equation for** *π* The update equation *q*(***π***) can be derived as follows.

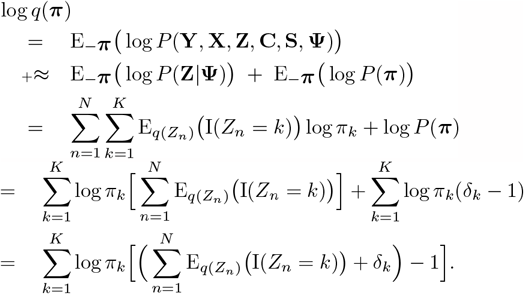

Therefore, *q*(***π***) is a Dirichlet distribution with parameters 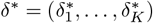, where

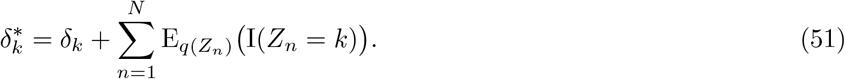

**Update equation for** *Z*_*n*_: The update equation *q*(*Z*_*n*_) can be obtained as follows. log *q*(*Z*_*n*_)

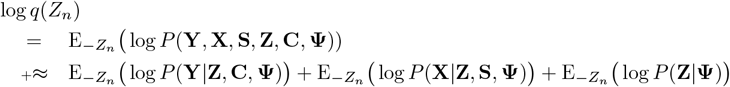

Note that, each of the logarithms above can be written as the sum of two terms, one that depends on *Z*_*n*_ and one that does not; since we want to form a function of *Z*_*n*_, w e discard all other term s w.r.t. *Z*_*c*_ : *c ∈* [*N*]\ *n*.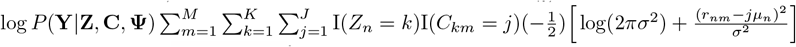 Taking the expectation, we have:

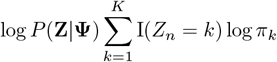

Combining the above parts from log *P*(**Y**|**Z, C, Ψ**) and log *P*(**Z**|**Ψ**), we obtain:

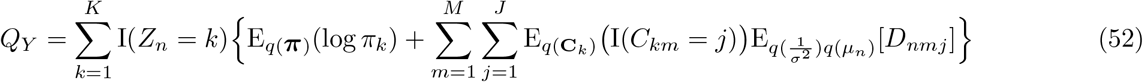

Where:

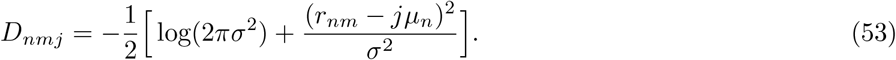

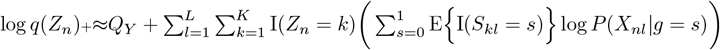

Reformulating log *q*(*Z*_*n*_), we achieve the below. Note that when processing the input data, for the missing values we merely calculate *Q*_*Y*_ .

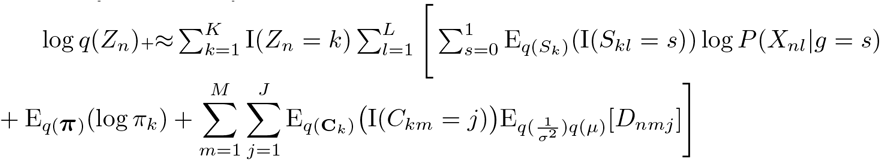

We conclude that *q*(*Z*_*n*_) *∼* Categorical(***p***_*n*_) with parameters ***p***_*n*_ = (*p*_*n*1_, …, *p*_*nK*_), where 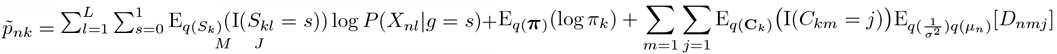 and

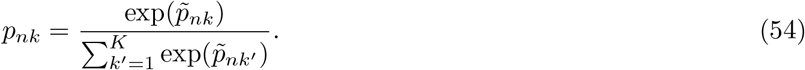

**Update equation for** ξ_*k*_: We calculate *q*(*ξ*_*k*_) as follows.

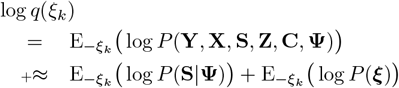

Disregarding the terms that do not depend on *ξ*_*k*_ in log *P*(**S**|**Ψ**) and log *P*(***ξ***), we obtain:

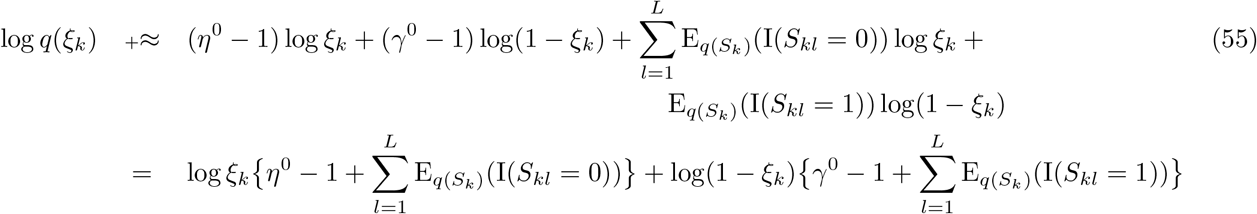

Therefore, *q*(*ξ*_*k*_) is distributed by Beta with parameters:

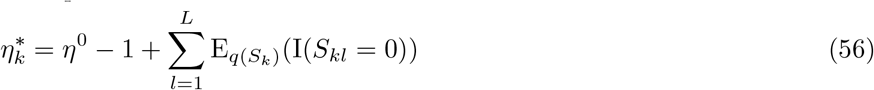

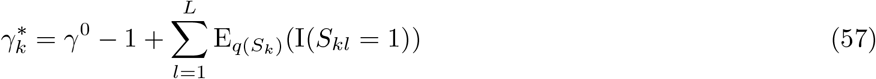

**Update equation for** *S*_*kl*_: We calculate *q*(*S*_*kl*_) as follows.

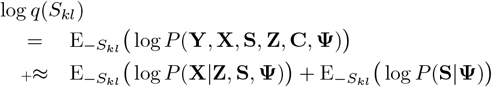

Disregarding the terms that do not depend on *S*_*kl*_ in the above, we obtain:

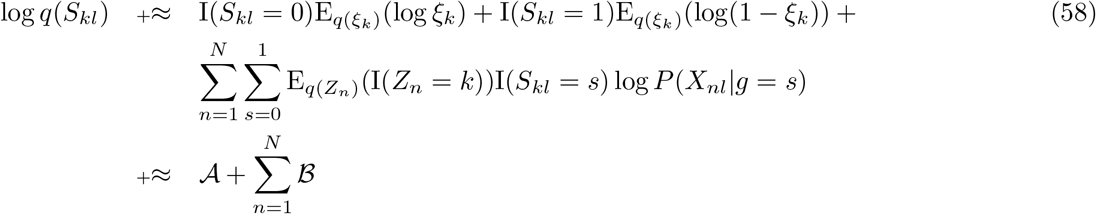

where 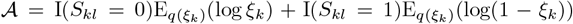, and 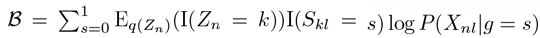.

Therefore, *q*(*S*_*kl*_) is distributed by Bernoulli up to an additive constant. The Bernoulli parameter, corresponding to *S*_*kl*_ = 1 is as the following. 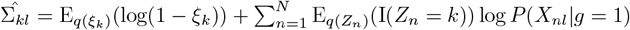

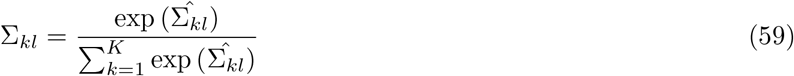

The rest of update equations are the same as those in CopyMix.

The new expectations that are used in the above update equations are the following.

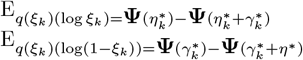

##### A.8 Simulation procedure

Here, we provide the details of the simulations. Because there is no simulation framework available, we generated data using the most reasonable way; that is, we assume a diploid cell (normal cell) can be affected by various copy number events (comprising duplications and deletions). We use an HMM, a generative model with latent copy number states, to generate the observed read ratios. HMM is a well-known model for copy numbers; see HMMCopy Shah et al. (2006) for more details.

For the Guassian emission, the generated configurations (CONF 1 to 18) are shown in the main paper. We modulate the read ratios by introducing subtle noise. Regarding the number of clusters, each of CONF 1 to CONF 5 contains two clusters, CONF 6 to CONF 10 three clusters, CONF 11 to CONF 13 four clusters, and CONF 14 to CONF 18 five clusters. The copy number transition patterns are formed using the following copy number state patterns, which are inspired by the CNV, along with single-cells that are obtained using the DLP technology Laks and McPherson (2019).

- **single state**: a copy number is inclined to remain at one certain state (CONF 1, CONF 2, CONF 3, CONF 4, CONF 5, CONF 12, and CONF 13 include this pattern);
- **inertial state**: a copy number is inclined to remain unchanged (The blue cluster in CONF 6 to CONF 10 includes this pattern);
- **oscillating state**: a copy number fluctuates between two states (The green cluster in CONF 6 and CONF 9 include this pattern);
- **altered state**: a sudden deletion-or duplication-like event of copy number occurs (all configurations include this pattern);
- **scaled state**: for specific ranges of positions in the copy number sequence, copy numbers are increased or decreased by a multiplicative or additive constant (all configurations include this pattern except CONF 3, CONF 4, CONF 6, CONF 11, and CONF 13);
- **whole-genome duplication**: all copy numbers across the genome are scaled (CONF 5 includes this pattern).

For both emissions, for each simulation, the following steps are performed.

1. Set the random seed.
2. Generate cells belonging to clusters by sampling from a multinomial distribution given a vector of cluster-assignment probabilities.
3. Generate rates for all cells. Rates are sampled from a Gaussian distribution with a mean of 10 and a standard deviation of 1.
4. Distribute the rates among the cells for different clusters; this is done based on step 2.
5. Set values to the number of HMM’s hidden states, transition matrix of each cluster, and sequence length.
6. Generate a Gaussian HMM for each cluster using cell rates of that cluster, number of hidden states, transition matrix of the cluster, and sequence length; this results in one copy number hidden sequence and different cell ratios emitted from that copy number sequence (the copy number is multiplied by the rate, representing the emission’s mean).
7. Accumulate the cells from the previous step and insert them into a dataset. Similarly, their cluster labels and the hidden copy number sequences are stored into datasets.

##### A.9 Ginkgo results

In this section, we report the results obtained by Ginkgo.

Ginkgo’s hierarchical clustering, after deleting 80% of the cells, detects three main clusters; firstly, the question is how the clustering would be if they didn’t exclude so many cells. Secondly, we cannot state that hierarchical clustering detects more than those three clusters. That is because the branch lengths of the smaller clusters, in the hierarchical clustering, are negligible in size compared to the branch length of the main three clusters. To include these branches and, consequently, the smaller clusters, one must choose a very shallow threshold when cutting the dendrogram produced by the hierarchical clustering.

If we, based on shared events in dark green in Fig. 10, assign a threshold such that we include the two subgroups depicted in the bottom and green part of Fig. 10, then Ginkgo detects four clusters. Regarding the four clusters, CopyMix outperforms Ginkgo with V-measure 67% compared to V-measure 55%. The remaining Ginkgo subgroups containing magenta-colored copy numbers require a lower threshold than those for the four clusters. Finally, it is essential to mention that Ginkgo’s web application, one advantage of using Ginkgo Mallory et al. (2020b), could not handle the large DLP data. To run Ginkgo, we developed a bash tool that assembled the back-end parts of the Ginkgo into a new script.

**Figure 10.**
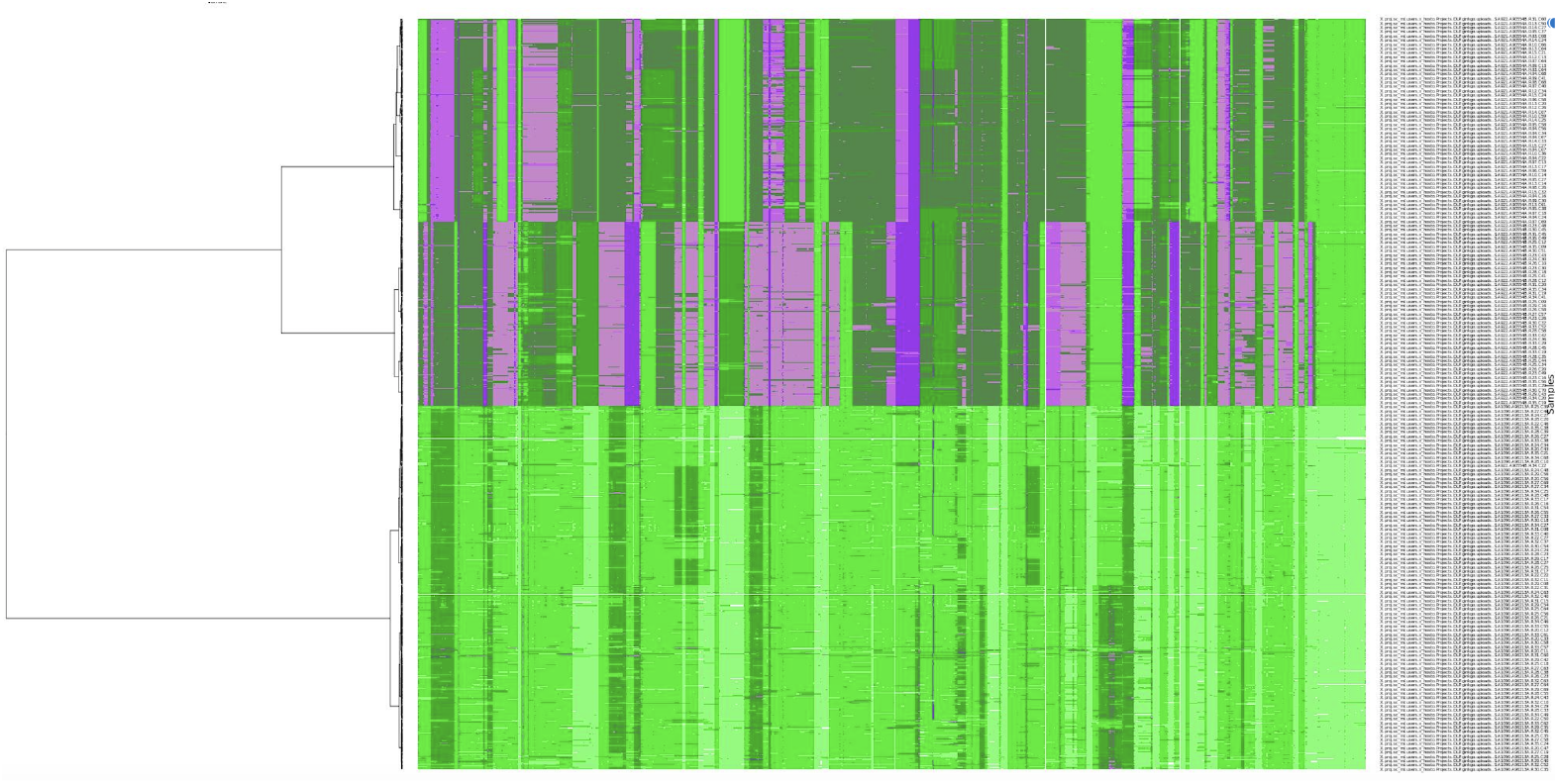
The heat map and the dendrogram show three major clusters detected by Ginkgo. The copy number is color-coded by a scale from green (small) to magenta (large).

##### A.10 Choice of Gaussian emission

This section discusses why Gaussian is chosen as a distribution for the read ratios. Following the principle of maximum entropy, we choose Gaussian because it has maximum entropy for a specified mean and variance Guiasu and Shenitzer (1985). Also, the histogram plot of DLP data (Fig. 11) confirms a skew-normal distribution. Finally, it is common that the data is assumed to follow Gaussian distribution; see HMMcopy and another HMM-based approach Shah et al. (2006); et al (2008).

**Figure 11.**
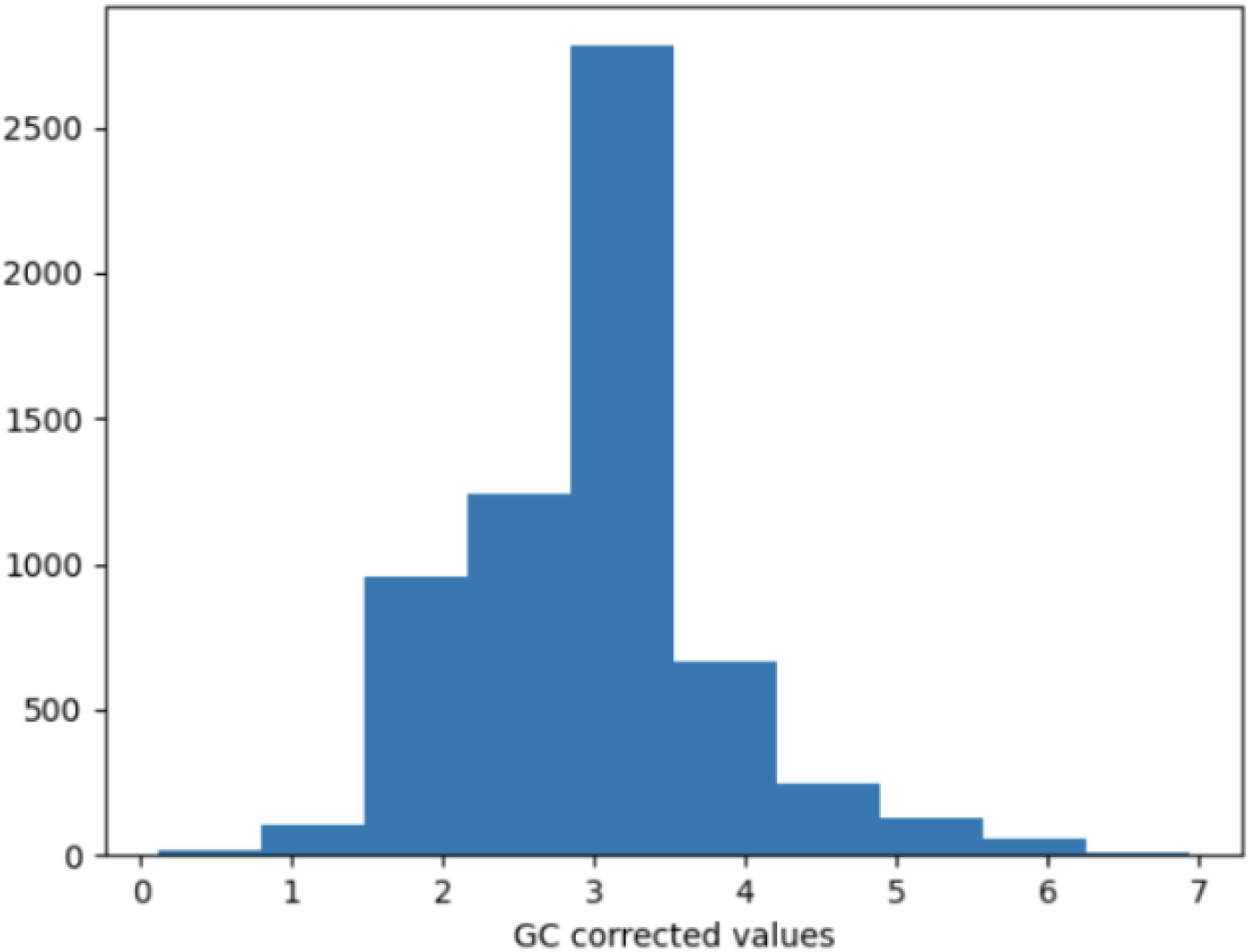
DLP data histogram plot.

##### A.11 SNV data

In this section, we illustrate the SNV data, confirming too shallow data signals for further cluster detection. As shown in Fig. 12, one can observe that the cells, which are on the Y axis, can be clustered into three main groups. Note that the three plots show in total 15000 sites where SNVs are detected. Performing K-means and hierarchical clustering resulted in three clusters; hence, it is reasonable that this data has been too shallow to reveal more clusters.

**Figure 12.**
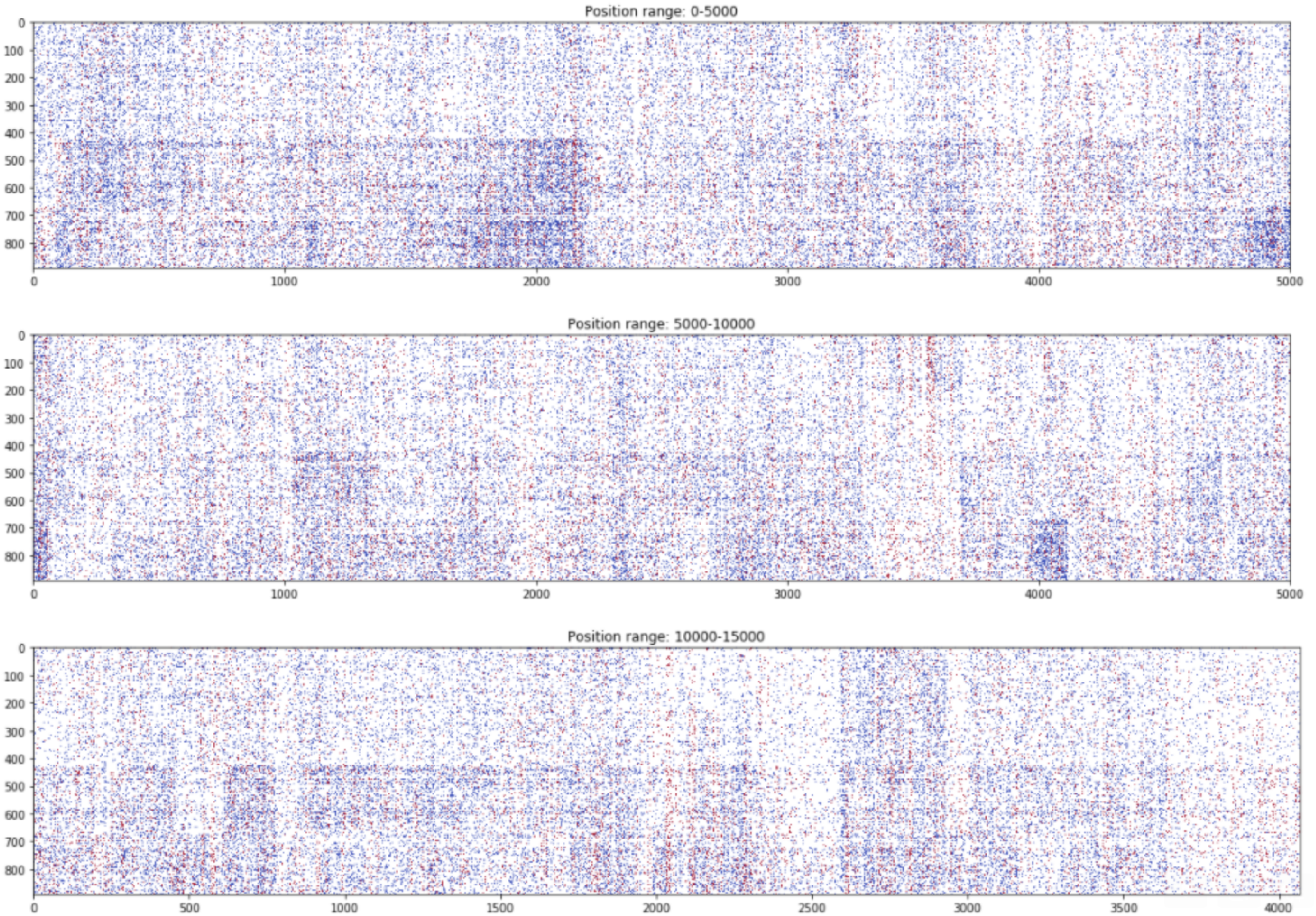
SNV counts are illustrated as colors in the heatmap for 891 cells (Y axis) across the genomic positions (X axis).

## Footnotes

3 CopyMix performance improves when the sequence is longer (the smaller the bin size, the longer the sequence) due to increasing the signal for the Markov chain. However, the low coverage in the DLP data increases the noise and missing data.

